# Mapping the peptide binding groove of MHC class I

**DOI:** 10.1101/2021.08.12.455998

**Authors:** Janine-Denise Kopicki, Ankur Saikia, Stephan Niebling, Christian Günther, Maria Garcia-Alai, Sebastian Springer, Charlotte Uetrecht

## Abstract

An essential element of adaptive immunity is the selective binding of peptide antigens by major histocompatibility complex (MHC) class I proteins and their presentation to cytotoxic T lymphocytes on the cell surface. Using native mass spectrometry, we here analyze the binding of peptides to an empty disulfide-stabilized HLA-A*02:01 molecule. This novel approach allows us to examine the binding properties of diverse peptides. The unique stability of our MHC class I even enables us to determine the binding affinity of complexes, which are suboptimally loaded with truncated or charge-reduced peptides. Notably, a unique erucamide adduct decouples affinity analysis from peptide identity alleviating issues usually attributed to clustering during electrospray ionization. We discovered that two anchor positions at the binding surface between MHC and peptide can be stabilized independently and further identify the contribution of other peptidic amino acids on the binding. We propose this as an alternative, likely universally applicable method to artificial prediction tools to estimate the binding strength of peptides to MHC class I complexes quickly and efficiently. This newly described MHC class I-peptide binding affinity quantitation represents a much needed orthogonal, independent approach to existing computational affinity predictions and has the potential to eliminate binding affinity biases and thus accelerate drug discovery in infectious diseases, autoimmunity, vaccine design, and cancer immunotherapy.

## INTRODUCTION

Major histocompatibility complex (MHC) class I molecules are central to the cellular adaptive immune response, presenting the intracellular peptidome to cytotoxic T lymphocytes, which detect non-self-peptide/MHC class I (pMHC) combinations and induce apoptosis in the presenting cell. Novel immunotherapy approaches, in which antiviral or antitumor T cells are identified, stimulated, and reintroduced into the patient, require the identification of high-affinity virus- or tumor-specific epitopes. The identification of high-affinity peptides is also key to developing efficacious peptide vaccines that have been prioritized by the WHO several years ago^1^. This is currently done either by isolating pMHCs from patient samples, eluting the peptides, and sequencing them by mass spectrometry (MS)^2^; or by sequencing the virus or the exome of a tumor and predicting the binding peptides^3^.

Such prediction is informed by the well-understood structural basis of MHC-peptide binding^4^, with > 1500 human and > 400 murine crystal structures in the protein data bank (PDB^5^) at this time. The peptide binds into a groove that consists of a β sheet topped by two parallel α helices with its amino terminus contacting the A pocket at one end of the groove^6^ and the *C*-terminal carboxylate held by a network of hydrogen bonds at the other end. These interactions limit the length of a high-affinity peptide to 8-10 amino acids^7^. In addition, side chains of the peptide (usually two) bind into other pockets at the bottom of the groove; one of them is always the F pocket (which usually binds the hydrophobic side chain of the *C*-terminal amino acid). Together, these interactions define the peptide binding motif of a particular class I allotype (such as x<u>L</u>xxxxxx<u>V</u> for HLA-A*02:01). About 16,000 class I allotypes have been described to date, with nine of them found in > 75 % of the Caucasian and Asian population^8^.

This structural knowledge, together with extensive databases of eluted peptides, has been used to train computational methods for the prediction of peptide epitopes from tumors^9^. However, this approach suffers from the uncertainty created by matching the eluted peptide to one of the six present MHC class I allotypes in a human being. Hence, to accelerate biological testing, it is desirable to rank candidate peptides by their binding affinity using a direct approach. Yet, to date, simple peptide binding assays that are amenable to high-throughput screening have not been available because they require empty peptide-receptive class I molecules, which are conformationally unstable^7, 10^.

Recently, empty disulfide-stabilized class I molecules (dsMHC) have become available^11–13^ that have the same peptide- and T cell receptor-binding specificities and affinities as the wildtype. Due to their stability in the empty state and their rapid binding, these dsMHC lend themselves to high-throughput approaches. We have previously employed native mass spectrometry (MS) to analyze peptide-bound MHC class I complexes. Furthermore, we have recently shown using native MS that the dsMHC variant of the HLA-A*02:01 (dsA2) can indeed be freed from the di-peptide used to facilitate refolding. Moreover, the empty binding groove can bind an exogenous peptide with the resulting pMHC appearing as an additional signal in the mass spectrum^14^. Native MS simultaneously detects all different mass species present in solution while quaternary structures and non-covalent bonds can be preserved^15^. Even though in the gas phase, there is no longer an equilibrium of protein-ligand binding, previous studies have demonstrated that native MS is an effective tool to determine dissociation constants, as the method captures and reflects the equilibrium state of the solution allowing the determination of absolute ligand binding affinities^16–22^.

In this work, the use of native MS to assess the binding affinity of peptides to dsMHC is demonstrated and verified by nanoscale differential scanning fluorimetry (nDSF). Our native MS method is used to show the effects of truncations and modifications on the binding affinity of peptides, the binding of two truncated peptides to the same class I molecule, and to investigate the cooperativity between the A and F pockets in peptide binding. Finally, yet importantly, this approach can be utilized to systematically investigate MHC class I peptide binding as the method is perfectly suitable for high-throughput applications.

## RESULTS

### Peak intensity in native mass spectra reflects peptide-MHC binding affinity

Empty dsA2 consists of the disulfide-stabilized heavy chain (hc, HLA-A*02:01(Y84C/A139C)) and the light chain, beta-2 microglobulin (β_2_m). Bacterially expressed dsA2 hc and β_2_m is folded *in vitro* into the dsA2 complex with dipeptides (GM or GL) and purified by size-exclusion chromatography (SEC) as described^10, 14^. During SEC, the dipeptide is removed and becomes undetectable by native MS, resulting in an empty binding groove (**Figure 1A and B**). At a low acceleration voltage of 25 V, raw and deconvoluted spectra (**Figure 1C and D**) demonstrate a stable complex of hc and β_2_m in the absence of any peptide. Some minor in-source dissociation (ISD, < 5 %) occurs upon activation in the mass spectrometer, with hc and β_2_m detected individually. The remaining fraction of dsA2 carries a small molecule adduct (337 Da by tandem MS analysis), which is easily released at ≥ 50 V by collision-induced dissociation (CID). Since protein production and sample preparation use no substance of this mass, the 337 Da molecule hypothetically originates from laboratory plastic ware as reported^23–25^. Indeed, small molecule MS reveals several common contaminants found in plastic-stored solutions, with erucamide ((*Z*)-docos-13-enamide) strongly enhanced in dsA2 samples and thus identified as the adduct (**Figure S1**).

**Figure 1.**
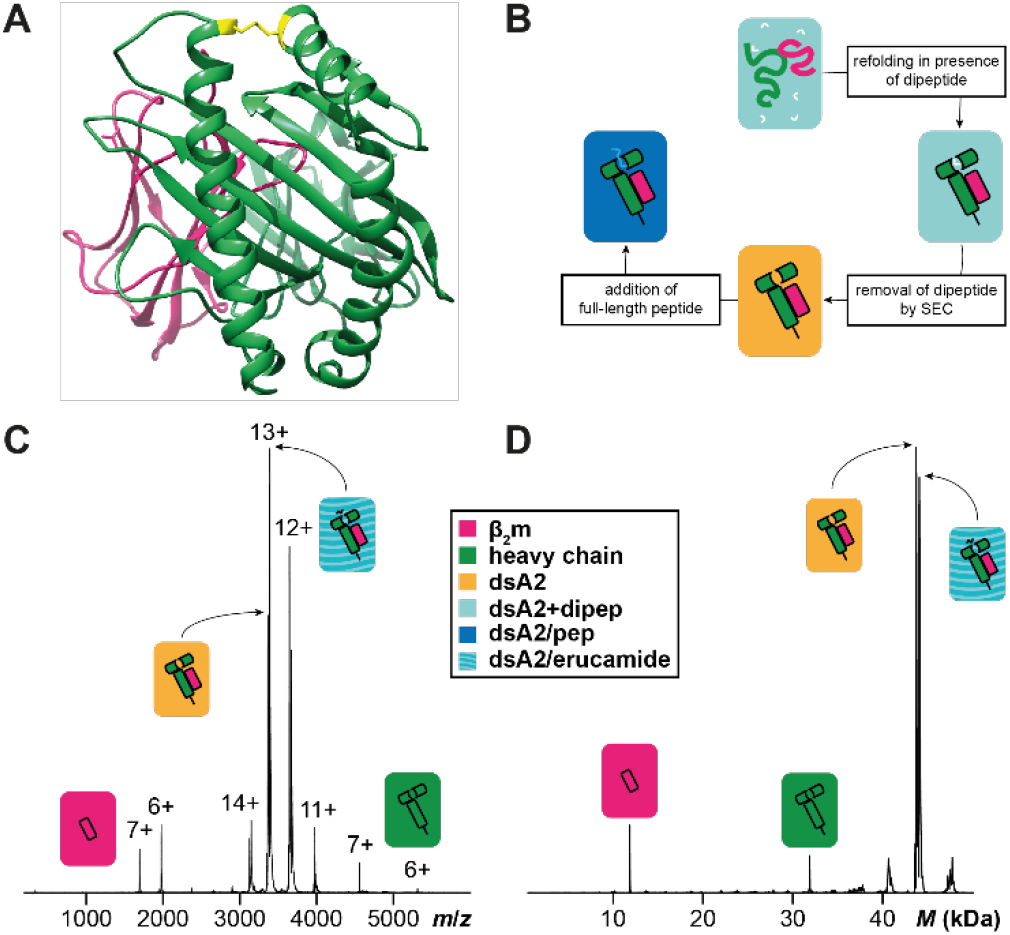
Disulfide-stabilized HLA-A*02:01. **(A)** Crystal structure of empty dsA2 (PDB 6TDR^14, 26^) seen as a top view of the peptide pocket. β_2_m is shown in pink, the heavy chain in green. The stabilizing disulfide bond (positions 84 and 139 mutated to cysteines) is depicted in yellow. **(B)** dsA2 is refolded in presence of GM or GL dipeptide, which is then removed via SEC. Afterwards, full-length peptides can bind into the empty binding groove. Raw **(C)** and deconvoluted **(D)** native mass spectra of peptide-free dsA2 recorded at an acceleration voltage of 25 V. The empty dsA2 is the predominant species (yellow). In addition, dsA2/erucamide (turquoise stripes) and dissociated β_2_m (pink) and heavy chain (green) can be seen.

In the following, it is examined whether native MS can differentiate the binding of high-affinity and low-affinity peptides by comparing the A2 epitope NV9 from human cytomegalovirus pp65 (sequence NLVPMVATV; theoretical dissociation constant, *K*_d,th_ = 26 nM predicted by NetMHC^9^) with the irrelevant YF9 (YPNVNIHNF; *K*_d,th_ = 27 μM) and with GV9 (GLGGGGGGV; *K*_d,th_ = 2.7 μM), a simplified NV9 derivative that retains the anchor residues, L and V. 10 μM dsA2 is incubated with 50 μM peptide for ten minutes prior to native MS, where the acceleration voltage across the collision cell is increased incrementally (10, 25, 50, 75, and 100 V) to observe the dissociation behavior. At 75 and 100 V, β_2_m heavily dissociates from the hc (data not shown), and so in the following, only results for 10 V, 25 V and 50 V are shown. The signal is quantified by determining the area under the curve (AUC) over the entire spectrum for each mass species (**Figure 2**; raw spectrum in **Figure S2**). While at 10 V and 25 V, the distribution of the different mass species is very similar, the ratios change significantly at 50 V. This is a frequent observation with the electrospray ionization (ESI) process, where non-covalent hydrophilic bonds such as those between dsA2 and high affinity peptides are retained^16, 27–29^, but hydrophobic interactions are weakened. By increasing the acceleration voltage, the dissociation of a protein-ligand complex usually does not occur gradually but spontaneously beyond a certain threshold, at which an energetic state is encountered that denatures the complex, in our case between 25 and 50 V. Therefore, the measurements at 10 V are used to calculate the dissociation constants.

**Figure 2.**
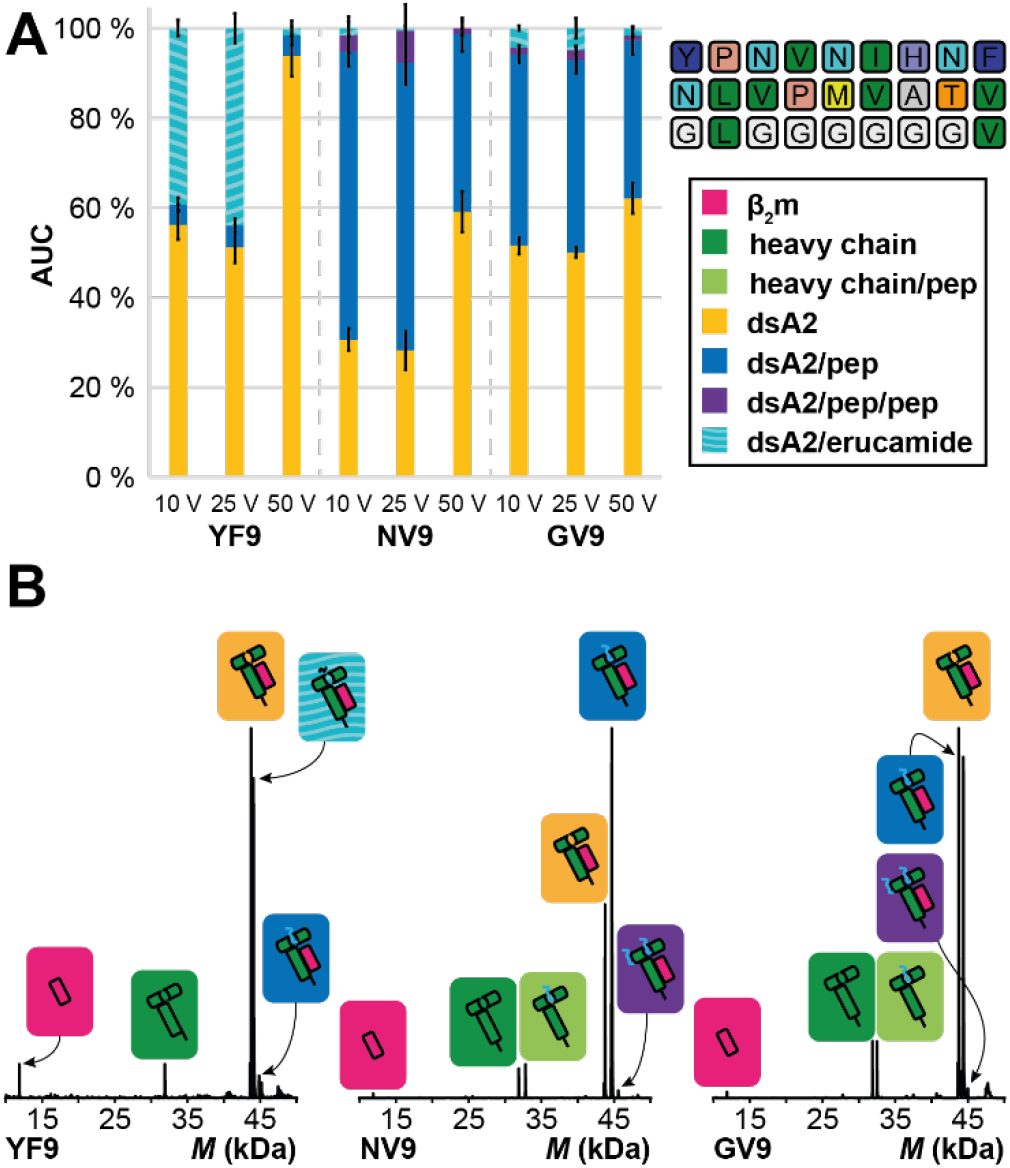
Overall area under the curve (AUC) for the detected dsA2 mass species in presence of YF9, NV9, and GV9 at different acceleration voltages. (**A**) The AUC is determined over the entire spectrum for the respective mass species at 10 V, 25 V, and 50 V. The mean value of the AUC in absence or presence of the different peptides (protein-peptide ratio 1:5) from at least three independent measurements is depicted along with error bars that represent the corresponding standard deviation. “dsA2” (yellow bars) corresponds to the empty HLA-A*02:01(Y84C/A139C) disulfide mutant complex, “dsA2/pep” (blue bars) to dsA2 bound to one peptide, “dsA2/pep/pep” to dsA2 bound to two molecules of this peptide (purple bars), and “dsA2/erucamide” to dsA2 bound to erucamide (turquoise-striped bars), respectively. The negative control YF9 barely associates with dsA2, whereas the positive control shows a high proportion of dsA2/NV9, indicating high affinity. GV9, which contains only the two anchor residues Leu-2 and Val-9 of NV9, still shows a high affinity, and at 50 V, their dsA2/pep proportions are very similar, showing that their gas phase stability is comparable. (**B**) Representative charge-deconvoluted spectra of the distinct protein and protein-peptide complex species recorded at 25 V. Pink and dark green correspond to the free β_2_m domain and heavy chain, respectively. Light green corresponds to a free heavy chain still attached to a peptide. The different complexes are assigned in yellow (empty dsA2), turquoise stripes (dsA2/erucamide), blue (dsA2/pep) and purple (dsA2/pep/pep).

In the presence of low-affinity YF9, the empty dsA2 molecule (43,733 ± 4 Da) generates the highest signal (56 ± 3 % at 10 V; **Figure 2A**). Most of the remainder of dsA2 carries only the erucamide adduct (44.071 ± 5 Da; 39 ± 2 %). There is very little dsA2/YF9 complex (4 ± 2 %), and because of the very low binding affinity of YF9, it can be assumed that this signal does not correspond to real binding events but rather to an artifact of the electrospray process known as non-specific clustering^17, 30–33^. Assuming that the other tested peptides cluster to the same extent, all native MS data is therefore corrected for the clustering determined with YF9. Since the data measured for the peptides of interest are therefore netted with YF9, no affinities are calculated for this control peptide. Corresponding raw data for the negative control are listed in the supplement (**Table S2**).

For NV9, in contrast, very efficient binding is observed, with 64 ± 3 % for dsA2/NV9 at 10 V and 40 ± 4 % at 50 V. Here, the dsA2/erucamide complex is completely absent, which suggests that erucamide is displaced by NV9. Erucamide either binds into the peptide groove, or it binds elsewhere and is displaced by a conformational change caused by peptide binding. A small amount of another mass species (45,624 ± 4 Da) that corresponds to dsA2 with two molecules of NV9 (dsA2/NV9/NV9) with an abundance of 4 ± 4 % (10 V) and 1 ± 2 % at 50 V is also observed. The latter is likely the result of unspecific clustering, as the abundance correlates with the intensity of the first binding event and is similar to the intensity of YF9 binding. Within NV9, the leucine in the second position and the *C*-terminal valine bind into the B and F pocket of the binding groove, respectively^12, 34–35^. The proportion of dsA2 occupied with GV9 at 10 V (43 ± 2 %, and 1.5 ± 0.4 % for dsA2/GV9/GV9) is significantly lower than the proportion of dsA2/NV9. This clearly shows that the minimal binding motif cannot support the same affinity as NV9, suggesting that other amino acids contribute significantly to the binding. At 50 V, however, the abundance of the dsA2/pep complex is the same for GV9 and NV9. This indicates that the strong B and F pocket side chain interactions together with the binding of the termini determine pMHC gas phase stability.

Despite very efficient binding, the obtained *K*_d_ for NV9 is only 8 ± 2 μM, while NV9 has nM affinity^36^. Hence, a fully occupied peptide binding pocket (predicted > 99 %) is expected in our measurements (**Figure 2A**). Protein denaturation due to storage or other physicochemical stress (data not shown) is excluded, and thus ISD, which occurs in the source region of the mass spectrometer and results in reduction of the dsA2/NV9 complex, is considered next. Here, lower cone voltages can increase the proportion of the occupied complex^37–38^. Indeed, the occupancy of the peptide binding groove with both the peptide and erucamide is higher at lower cone voltages. This linear relationship is most evident with dsA2 alone, to which erucamide is the only binding partner (**Figure 3A**). The data suggest that at a cone voltage of ≤ 36 V, which is unfeasible experimentally, dsA2 is 100 % occupied with erucamide. This observation allows us to use erucamide as a reference species for peptide binding measurements, assuming that in solution, all free dsA2 protein is initially bound to erucamide as suggested by the zero cone voltage extrapolation, and that the peptide replaces it in the binding region. From the erucamide-bound fraction measured at a cone voltage of 150 V in native MS, the fraction of dsA2/erucamide corresponding to the fraction of dsA2 not bound to peptide at the end of the binding reaction can be recalculated using the correction factor of 2.2 originating from the linear function’s slope. From this, the fraction of dsA2/peptide is inferred, resulting in an apparent *K*_d_ that reflects the in-solution environment (see the Materials and Methods). This is highly advantageous, since for individual pMHCs, the ISD is naturally influenced by peptide size and sequence, precluding compensation, while the ISD of the MHC-erucamide complex is invariant and peptide-independent. Therefore, for each peptide both a dissociation constant for the high cone voltage (150 V) based on clustering-corrected MHC-peptide signal (*K*_d,high_) and another for the theoretical low cone voltage of 36 V based on the MHC-erucamide signal (*K*_d,low_) is described (**Figure 3A**). While the experimentally determined *K*_d,high_ is overestimated due to ISD, the *K*_d,low_ is a more reliable approximation.

**Figure 3.**
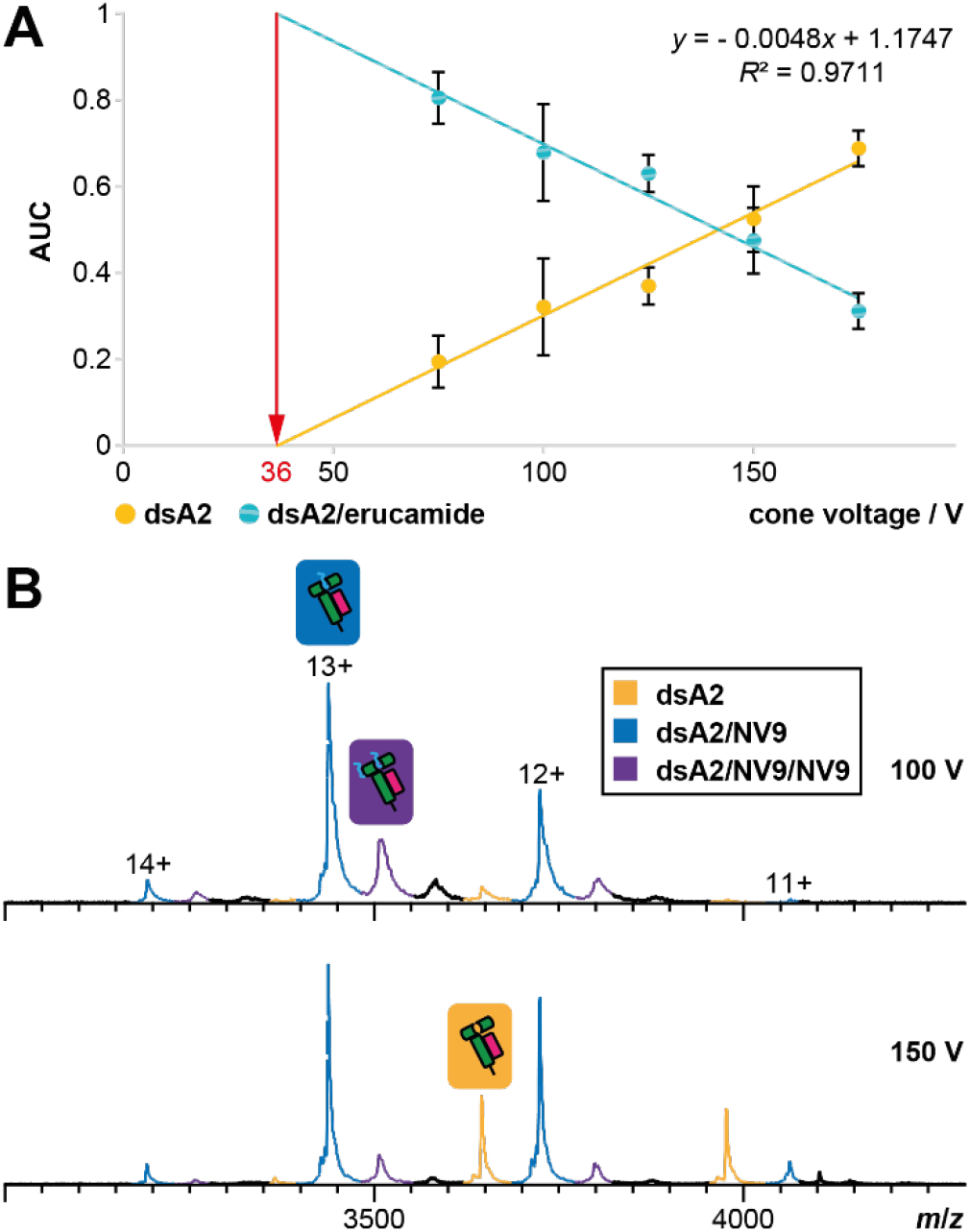
Ion-source decay causes linear dependence of the ligand-bound dsA2 fraction on cone voltage. (A) Native MS fractions of dsA2 and dsA2/erucamide are likewise dependent on the activation energy of the cone voltage. As the cone voltage is raised, the percentage of empty dsA2 increases because the adduct binding cannot withstand the energy in the ion source and therefore dissociates. The proportionality is linear, and therefore a theoretical cone voltage of 36 V can be calculated at which a full occupancy with ligand would be reached. In reality, this voltage is too low to obtain stable electrospray and a resolved spectrum. (B) Peptide binding is equally subjected to in-source dissociation. NV9-bound dsA2 (blue and purple annotated peaks) become less prominent with increasing cone voltage, while empty dsA2 (yellow) increases at the same time.

To see whether the affinities thus obtained match those measured by a previously validated method, a series of sixteen peptides, most of them variations or truncations of the sequence of NV9, are compared in native MS and in isothermal analysis of nanoscale differential scanning fluorimetry (iDSF)^39^. *K_d_* values from native MS and iDSF correlate very well over a wide range (**Table S3**; **Figure 4A**). It has to be noted though that affinities below 200 nM have to be treated with caution as the peptide concentration, and hence the fraction of dsA2/pep, is very low. This is also evident from the thermal stabilization discussed in the following. For NV9, the binding groove is so well stabilized that other domains unfold first^40–41^. Nevertheless, the peptide binding affinity measured by native MS correlates very well with the thermal stabilization of dsA2 by peptide binding (protein-peptide ratio: 1:10, **Figure 4B**). Conformational stabilization corresponds to an increase in the midpoint of thermal denaturation (*T_m_*) above that of the empty dsA2, which is measured by tryptophan nanoscale differential scanning fluorimetry (nDSF^40^, ^42^) to be 35.7 ± 0.6 °C. While the negative control YF9 clearly shows no (*ΔT_m_* = 1 ± 1 K), the positive control NV9 shows a high degree of stabilization (*ΔT_m_* = 23.4 ± 0.6 K) in agreement with published data^12^. The other peptides show excellent correlation between affinity and stabilization ability. All peptides identified as strong binders by their apparent dissociation constant exhibit a *ΔT_m_* of at least 6.4 ± 0.6 K (GV9), while the *ΔT_m_* for the low-affinity peptides ranges between ≈ 0 K and ≈ 3 K (**Figure 4B**, red area). Closer evaluation of the data shows that peptides with a *K*_d,low_ < 1 μM also have an increased gas phase stability of the complex at 50 V (dsA2/pep + dsA2/pep/pep > 35 %) and present only negligible amounts of dsA2/erucamide at any voltage. Therefore, we define all peptides that fall within this range as strong binders for dsA2 (**Figure 4B**, white area). These results demonstrate that by simply comparing the intensities in the native mass spectrum, it is possible to estimate, which properties define peptides to act as high-affinity epitopes for HLA-A*02:01.

**Figure 4.**
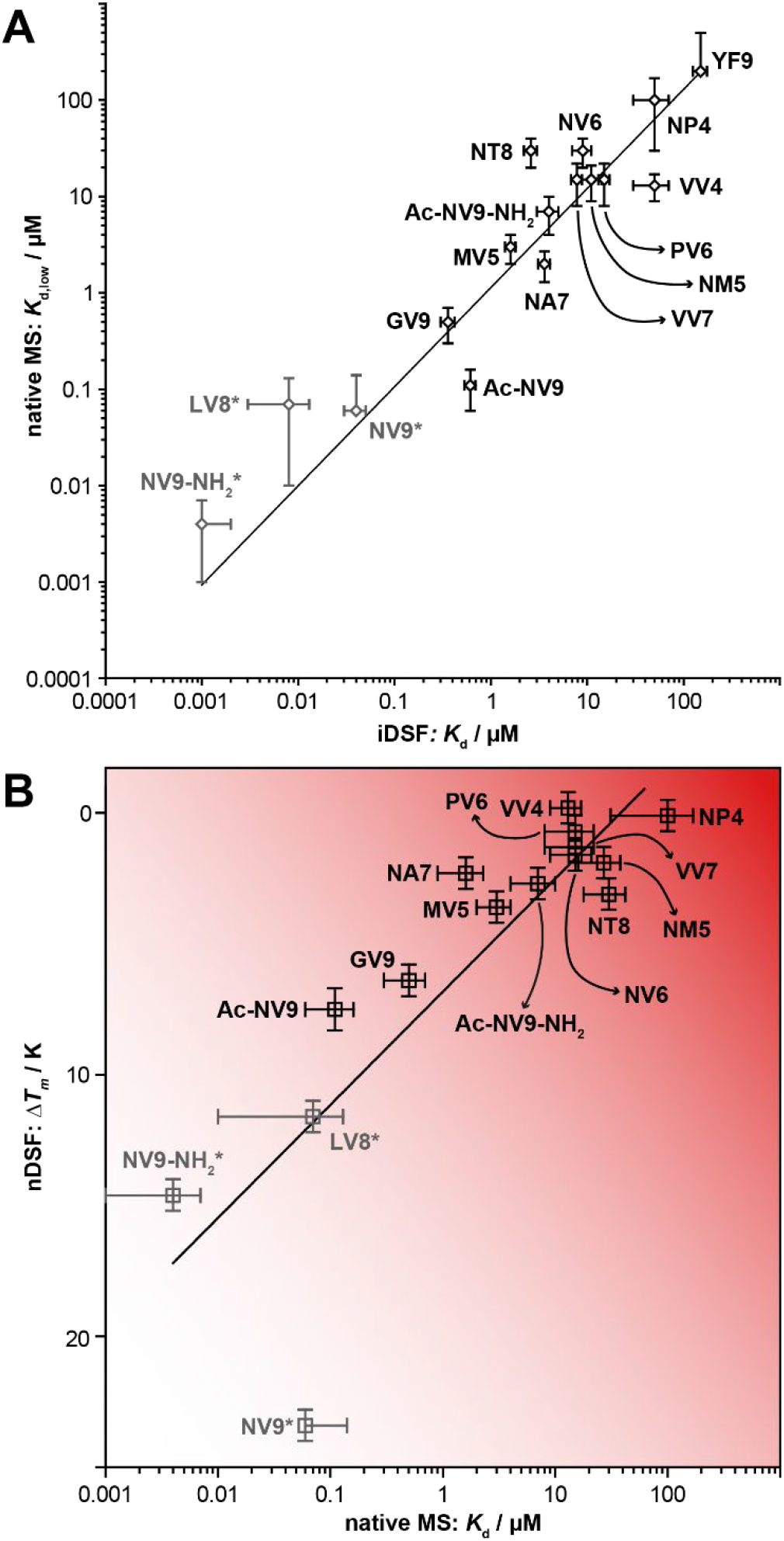
Relation of thermal stability and affinity for dsA2/peptide complexes. **(A)** Affinities determined via iDSF and native MS correlate. Displayed data points represent the relationship of both apparent *K*_d_ for all different dsA2/peptide systems analyzed. Both axes are scaled logarithmically. **(B)** Thermal denaturation measurements availing intrinsic tryptophans’ change in fluorescence are used to define protein complex stabilization upon peptide binding, whereas an apparent *K*_d_ for the various peptides is determined using native MS (10 V acceleration voltage, 150 V cone voltage). The dsA2 complex and peptide are deployed in a ratio of 1:10 (*ΔT_m_*) or 1:5 (*K*_d_) depending on the experiment. Peptides showing a small *K*_d,low_ for binding dsA2 and being concomitantly able to stabilize the protein complex are defined as strong binders (white area). The remaining peptides (red area) lack crucial features, making them unable to form strong bonds to dsA2 indicated by low binding affinities and melting temperatures. Standard deviation for both methods is displayed by error bars. The x coordinate is displayed logarithmically. *iDSF reaches its limits at affinities below 200 nm, hence the values of grayed-out peptides are not reliable.

### Neutralizing the terminal charges of the peptide reduces binding efficiency

Next, the influence of the charged termini of the peptide upon the binding affinity is analyzed. For this purpose, three variants of NV9 are designed: Ac-NV9-NH_2_ has an acetylated *N*-terminus and an amidated *C*-terminus, whereas Ac-NV9 and NV9-NH_2_ each carry only one of these modifications. For the dsA2/pep fraction at 10 V, Ac-NV9 and NV9-NH_2_ show only a small difference to the unmodified NV9, resulting in comparable apparent *K*_d_ values (*K*_d,low_ = 0.11 ± 0.05 μM and 0.004 ± 0.003 μM). Further, no increase of dsA2/erucamide is observed (**Figure 5A**). For both peptides, the protein-ligand complex is still stable at 50 V (**Figure S5**). For Ac-NV9, the proportion of the occupancy is even higher than for NV9 itself (57 ± 2 % and 8 ± 1 % vs. 40 ± 4 % and 1 ± 2 %, **Table S2**). Remarkably, all modified peptides have an increased double occupancy. For the previously discussed peptides, dsA2/pep/pep is significantly lower with the result that correction for non-specific clustering reduces it to a value below threshold. For Ac-NV9, this effect is significant, since even at 50 V the proportion of dsA2/pep/pep (purple bars) is still 8 ± 1 %. However, the stabilization effect on dsA2 in a 1:10 thermal denaturation approach is rather moderate for Ac-NV9 (*ΔT_m_* = 7.5 ± 0.8 K), while it is very strong for NV9-NH_2_ (*ΔT_m_* = 14.6 ± 0.6 K), indicating that the *N*-terminus has more relevance for tight peptide binding than the *C*-terminus.

**Figure 5.**
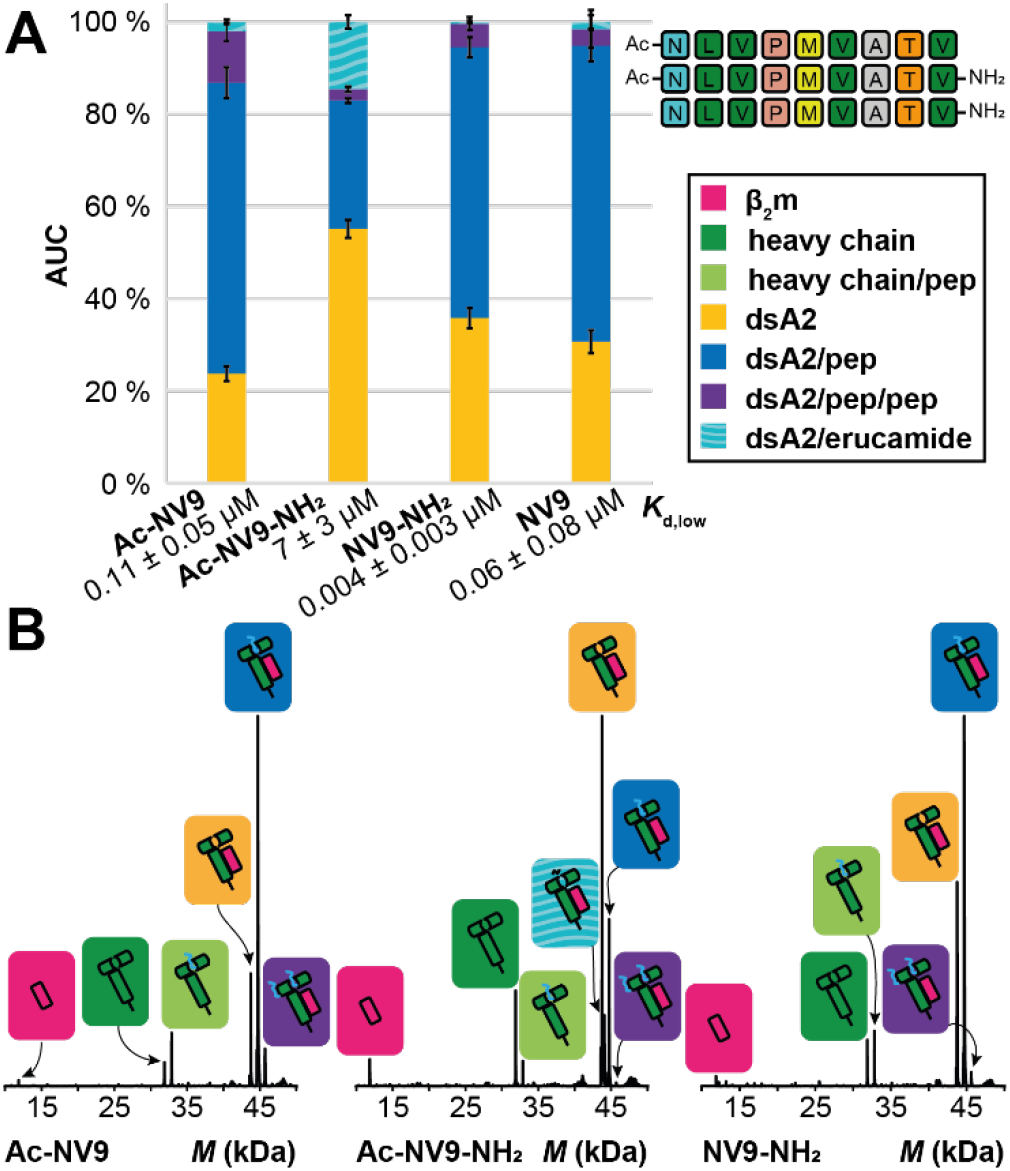
Charge-reduced NV9 variants analyzed at 10 V. **(A)** The AUC is determined over the entire spectrum for the respective mass species at an acceleration voltage of 10 V. The mean value of the AUC in absence or presence of the different peptides (protein-peptide ratio 1:5) from at least three independent measurements is depicted along with error bars that represent the corresponding standard deviation. “dsA2” (yellow bars) corresponds to the empty complex, “dsA2/pep” (blue bars) to dsA2 bound to one peptide, “dsA2/pep/pep” to dsA2 bound to two molecules of this certain peptide (purple bars) and “dsA2/erucamide” to dsA2 bound to erucamide (turquoise-striped bars) respectively. By modifying only one terminus (Ac-NV9 and NV9-NH_2_), the affinity of the peptide to dsA2 changes only marginally, but if the charges on both termini are neutralized (Ac-NV9-NH_2_), the peptide binding is greatly reduced. The corresponding *K*_d,low_ is shown for respective peptides. **(B)** Representative charge deconvoluted spectra of the distinct protein and protein-peptide complex species recorded at 10 V. Pink and dark green correspond to the free β_2_m domain and heavy chain respectively. Light green corresponds to a free heavy chain still attached to a peptide. The different complexes are assigned in yellow (empty dsA2), turquoise stripes (dsA2/erucamide), blue (dsA2/pep) and purple (dsA2/pep/pep).

The correction for unspecific binding is based on the negative control YF9, with both termini unmodified. By neutralizing the positive charge at the amino group, Ac-NV9 could be more susceptible to nonspecific electrostatic interactions at the protein surface, which are then preserved in the gas phase. However, this should not be the case for the erucamide-based *K*_d_. The variant Ac-NV9-NH_2_ shows significant changes both in its affinity to dsA2 and in complex stabilization. At the same time, there is no significant increase in the formation of double binding events. The affinity decreases significantly (*K*_d,low_ = 7 ± 3 μM) due to the dual modification, a high amount of erucamide is not replaced, and at an acceleration voltage of 50 V, the complex is no longer stable. The nDSF measurements likewise show that the stabilization effect with *ΔT_m_* = 2.7 ± 0.6 K is rather weak. Although NV9-NH_2_ is far better rated within our scheme (**Figure 4A**) in terms of affinity and stabilizing effect, Ac-NV9 still falls into the category of a strong binder. For the dual-modified variant, which no longer carries terminal charges, a clear loss of binding efficiency can be observed indicating that ionic interactions are indispensable in the formation of the MHC-peptide complex and, without them, efficient binding cannot be established.

### Truncated NV9 discloses preferred binding positions in the peptide

In the following, it is analyzed how stepwise truncation of NV9 from either terminus affects binding to identify the binding contributions of individual residues by MS. Building on the knowledge from the dsA2/NV9 crystal structure^12, 14^, the peptide binding groove can thus be mapped and further understood. *In vivo*, MHC class I predominantly binds peptides with a length of eight to ten amino acids^7, 43–44^. It is therefore not surprising that octa- and nonapeptides show the highest affinity (**Figure 6, Table S3**). Looking at the effect of the loss of only one of the two terminal amino acids, asparagine and valine, the difference is striking. Despite the absence of asparagine (corresponding to LV8), the affinity (*K*_d,low_ = 0.07 ± 0.06 μM) is still comparable to NV9 with a considerable stabilizing effect during thermal denaturation of dsA2 (1:10, *ΔT_m_* = 11.6 ± 0.6 K). At 50 V, dsA2/LV8 even appears to be more stable than dsA2/NV9. In contrast, loss of the *C*-terminal valine results in a strong decrease in affinity (*K*_d,low_ = 30 ± 10 μM). The erucamide is no longer fully displaced by NT8, only a small fraction of the protein-peptide complex is resistant to the higher acceleration voltage and no significant stabilization effect (*ΔT_m_* = 3.1 ± 0.6 K) is measured. These observations suggest that the binding between the *C*-terminus of the peptide and the F pocket of the MHC molecule is essential, whereas the binding of the *N*-terminus located within the A pocket is less important for the overall binding efficiency. However, when the second amino acid in the *N*-terminal direction, leucine, is eliminated as well (VV7), affinity and stabilizing effects are likewise diminished. Along with our findings on GV9, it is evident that the anchor residues of the nonapeptide are in second and last position. Efficient binding between peptide and HLA-A*02:01 only occurs if both positions are covered by suitable amino acids. Nevertheless, the difference between GV9 and LV8 points towards an additional contribution of the other amino acids that is in sum larger than the *N*-terminal contribution.

**Figure 6.**
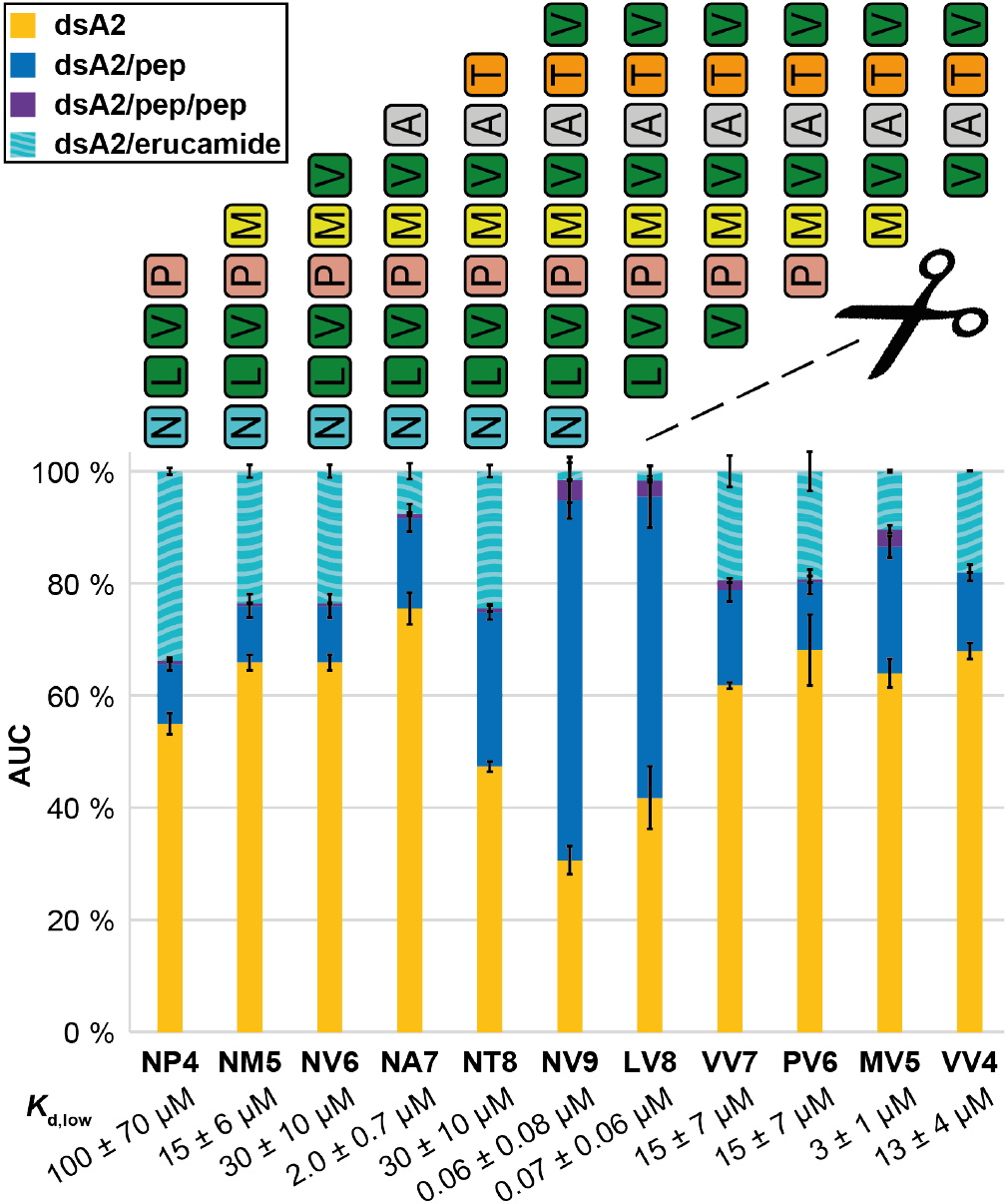
Truncated NV9 variants analyzed at 10 V. The AUC is determined over the entire spectrum for the respective mass species at 10 V. The mean value of the AUC in absence or presence of the different peptides (protein-peptide ratio 1:5) from at least three independent measurements is depicted along with error bars that represent the corresponding standard deviation. “dsA2” (yellow bars) corresponds to the empty complex, “dsA2/pep” (blue bars) to dsA2 bound to one peptide, “dsA2/pep/pep” to dsA2 bound to two molecules of this certain peptide (purple bars) and “dsA2/erucamide” to dsA2 bound to the erucamide (turquoise-striped bars) respectively. Octa- and nonapeptides have the highest binding efficiency in accordance with their biological function. Without the anchor residues Leu-2 and Val-9 in a truncated NV9 variant, the affinity decreases drastically. A terminal methionine appears to enhance the affinity of the peptide to dsA2. The corresponding *K*_d,low_ is shown for respective peptides.

Therefore, none of the examined tetra-to heptapeptides are strong binders. In direct comparison, VV7, which still contains the *C*-terminal anchor residue Val-9, performs worse than NA7, which in contrast carries Leu-2. NA7 displaces more of the erucamide, and at 50 V, significantly more NA7 than VV7 is observed (**Figure S5**). Thus, Leu-2 appears to possess more binding strength than Val-9. Curiously enough, the *N*-terminally truncated pentapeptide MV5 performs significantly better than the corresponding hexa-(PV6) and heptapeptides (VV7). The increased proportion of dsA2/MV5, which is even more evident at 50 V (**Figure S5**) and the simultaneous occurrence of dsA2/MV5/MV5 (purple bars, **Figure 6**) strongly suggest that this peptide occupies an additional binding site within the peptide groove. Since this effect is not apparent for the smaller and therefore less spatially restricted VV4, the terminal methionine seems to be the decisive factor here. Since no increased dual occupancy is observed for NM5 and since binding at 50 V is weaker in direct comparison to MV5, it seems that the terminal methionine as such is not determinant on its own, but much more the relative position within the peptide. Considering the tetrapeptides, again the *N*-terminally truncated peptide, VV4, performs better than the *C*-terminally truncated NP4 at all acceleration voltages, which points to the positioning of the erucamide within the peptide pocket. While NP4 carries only the anchor peptide Leu-2, it seems that VV4 has a higher chance to bind to the dsA2 peptide groove due to its two terminal valines.

### The A and F pocket may be occupied simultaneously by two peptides

Encouraged by MV5 double binding, the simultaneous presence of two peptides in the binding pocket is further investigated^45^. Since binding of either peptide can be measured independently of the presence of the other in a single experiment – which, so far, no other method can provide – this allows us to assess whether the binding to the two ends of the peptide binding groove of MHC class I is cooperative. Our experiments are done with the partial peptides of NV9 that are combined with their corresponding counterparts from the opposite terminus. There are two ways in which two short peptides that bind to different sites in the binding groove could synergize in binding. They may stabilize the conformation of the molecule independently of each other, without any communication between the binding sites. In addition, it is possible that they bind in a cooperative manner. Hereby, the two binding sites communicate by a conformational or dynamic change in the protein and one peptide stabilizes (or destabilizes) the binding site for the other, such that the binding affinity for the second peptide is higher (or lower) in presence of the first.

In **Figure 7A**, single and double occupancy for the first and second peptide as well as the double occupancy of both peptides are depicted. The results of the native MS measurements reveal that the previously presented affinities of the two individual peptides are also reflected well in this combined setup. This also implies that the overall proportion of dsA2/pep1/pep2 remains very low in absence of positive cooperativity. Presumably, due to spatial restriction, no double-occupied complex can be detected in the pairing of the pentapeptide NM5 and the hexapeptide PV6. In the other combinations, it is possible to detect dsA2/pep1/pep2. Yet only when MV5 is involved in the binding, the complex is able to withstand higher acceleration voltages. It is assumed, however, that this is not due to a cooperative binding between NP4 and MV5, NM5 and MV5 or NV6 and MV5, but can rather be explained by the higher affinity of MV5, which again emerges in this experiment. With the dual approach, it is visible that VV4 is more dominant than NP4. The indirect comparison between NP4+PV6 and NV6+VV4 highlights this in particular. The shorter VV4 achieves a significantly higher occupancy fraction in competition with the corresponding hexapeptide, which is not the case for NP4. This once again suggests that VATV appears to have more binding options due to its two terminal valines.

**Figure 7.**
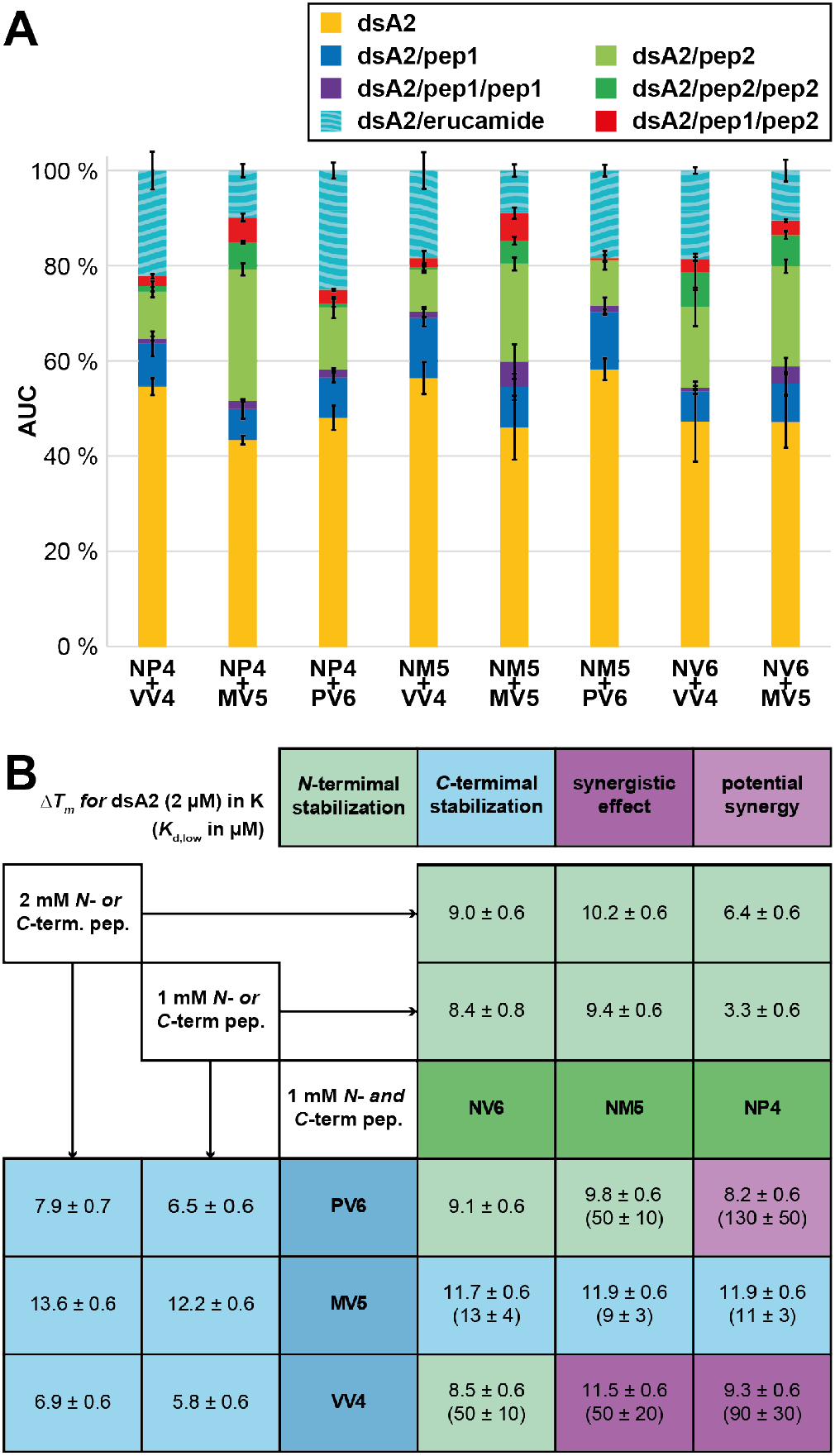
Detected dsA2 mass species in presence of two corresponding truncated NV9 variants. **(A)** The native MS data suggest that the truncated peptides do not bind cooperatively as the amount of dsA2/pep1/pep2 remains small in all measurements. Rather, the affinity of the individual peptides is independent from each other. The AUC is determined over the entire spectrum for the respective mass species at 10 V. The mean value of the AUC in absence or presence of the different peptides (protein-peptide ratio 1:10:10) from at least three independent measurements is depicted along with error bars that represent the corresponding standard deviation. “dsA2” (yellow bars) corresponds to the empty HLA-A*02:01(Y84C/A139C) disulfide mutant complex, “dsA2/pep” (blue bars) to dsA2 bound to one peptide, “dsA2/pep/pep” to dsA2 bound to two molecules of this certain peptide (purple bars), “dsA2/pep2” to dsA2 bound to another peptide when two different peptides where present (light green), “dsA2/pep2/pep2” to dsA2 bound to two molecules of the second peptide (dark green), “dsA2/pep1/pep2” to dsA2 bound to one molecule of each of both peptides (red bars) and “dsA2/erucamide” to dsA2 bound to the erucamide (turquoise-striped bars) respectively. The corresponding *K*_d,low_ is shown for respective peptide pairs. **(B)** The matrix visualizes the changes of *T_m_* in case half of the peptide amount is either exchanged for the corresponding *N*-(green) or *C*-terminal peptide respectively or alternatively omitted. Light green marks the occurrence of major *N*-terminal stabilization whereas for the light blue cells, the complex is primarily stabilized by the *C*-terminal peptide. Purple cells show peptide coupling with significant synergistic effect or potential synergy (light purple).

While in the native MS measurements, the occupancy of two peptides can be observed simultaneously, in nDSF we can use higher peptide excess and evaluate whether two peptides are able to synergistically stabilize the complex. The melting temperature of dsA2 exposed to only one of the truncated NV9 variants (**Figure S3**) is compared to the resulting *T_m_* when two peptides are utilized. The ratio between protein and total peptide concentration is kept constant at 1:1000. Assuming no synergistic effect occurs, the resulting melting temperature is expected to be just between those for the individual competing peptides according to their affinity to dsA2. The majority of the peptide pairs show a competitive behavior and do not demonstrate better stabilization by both peptides combined. Whereas for NP4 and VV4 and to a lesser extent for NP4 and PV6 as well, a higher *T_m_* is measured for both peptides together than for each of them separately revealing that dsA2 can be stabilized by two shorter peptides in a synergistic fashion while the respective affinities are not mutually influenced (**Figure 7B**). Once again, MV5 stands out particularly in this experiment. Two parts of MV5 provide significantly more thermal stability to dsA2 than one part of MV5 together with one part of one of either of the corresponding *N*-terminal peptides. These observations suggest that unfolding of MHC class I or at least of the disulfide mutant can begin at either end of the peptide binding groove. Consequently, this also means that the opposite ends of the binding groove cannot communicate conformationally, which was already described for the corresponding wildtype^11^.

## DISCUSSION

While only wildtype pMHC were studied by native MS so far^46^, we have recently demonstrated that peptides added to empty disulfide-stabilized class I molecules can be detected as well^14^. This study shows that this method, which is fast and amenable to high-throughput approaches, can be used to measure MHC class I peptide binding affinities, and it is used to map the contributions of parts of the peptide to high-affinity binding.

Our data affirms the key interactions between dsA2 and its high-affinity ligand NV9 that were described previously^47–49^. It is therefore scarcely surprising that Leu-2 and the *C*-terminal Val-9 contribute the major portion of the total binding. Leucine binds in the B pocket and valine in the F pocket of the binding groove, respectively. The centrally located Val-6 appears to play a minor role in binding, as long as full-length NV9 is considered. However, when situated in second position due to truncation, valine can act as an anchor residue as well. For the F pocket, it is well established that Val or Leu are the strongest anchor residues, but our data suggest that MVATV can also stabilize the B pocket via its first valine similarly and possibly even simultaneously. The extraordinary thermal stabilization observed for dsA2/MV5, especially in comparison with potentially synergistic peptide pairs, gives rise to the assumption that even more than two binding modes are available for MV5. As reported before^50^, along with leucine and methionine, valine is indeed one of the amino acid residues preferred by the A*02:01 and moreover by the entire A2 supertype as an anchor residue in second position. Since proline in the *N*-terminal position was found to be a deleterious factor for binding, this could explain the poor performance of PV6 in our experiments despite the preferred methionine in the second position. Surprisingly, NV6 scores low in all our tests despite having leucine in the second position and a *C*-terminal valine. Given the two different binding modes, at least a higher proportion of singly bound species would be expected here – similar to MV5 – for statistical reasons. A potential explanation may be that the peptide competes with itself for the two separate binding sites. Based on their sheer size, two NV6 molecules sterically hinder each other as opposed to the pentapeptide MV5. As previously demonstrated^14^, the F pocket can be occupied by GM with methionine as anchor residue. Methionine in the *C*-terminal position is not preferred but only tolerated by HLA-A*02:01, as reported before^50^. Compared to MV5, which can bind with either valines, the second pentapeptide, NM5, therefore shows the same strong binding behavior in our experiments. For larger peptides like NV9, which were found to be double bound to dsA2, it is more likely that the signal originates from nonspecific clustering of NV9 with the abundant dsA2/NV9 complex rather than from the actual binding of two peptides to the dsA2 peptide binding groove due to spatial reasons.

Parker *et al*.^48^ claimed that GLGGGGGGV, carrying the minimal binding motif, is not sufficient for stabilization of HLA-A*02:01. However, GV9 is a strong binding ligand in our native MS experiment, and GV9 binding also increases thermal stability. Nevertheless, there are significant differences compared to the high-affinity control NV9. Hence, it is clear that other amino acids contribute substantially to overall binding. We hereby propose that the disulfide-stabilized MHC class I molecule has at least two positions that can be stabilized independently upon peptide binding as seen for the shorter peptides. **Figure 8** presents the binding possibilities within the HLA-A*02:01 peptide pocket identified in this study.

**Figure 8.**
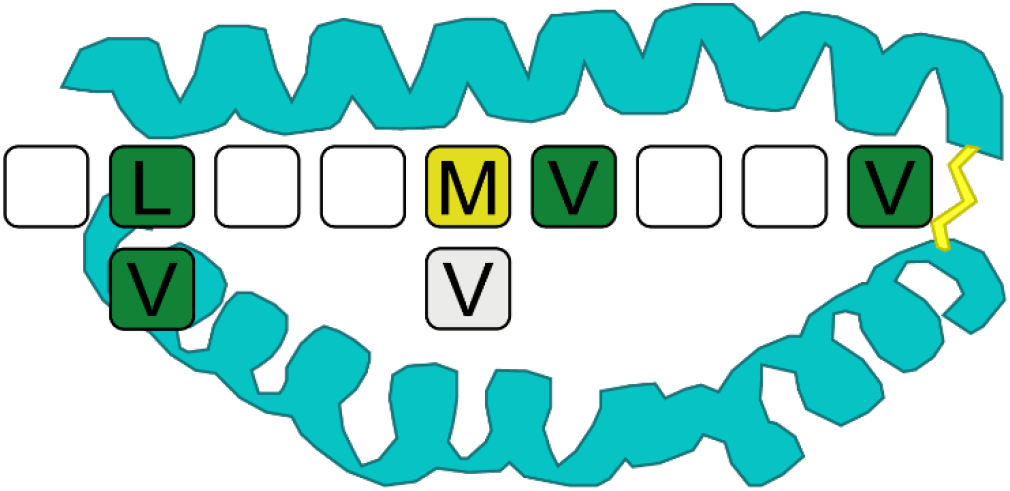
Favored amino acid positions within the HLA-A*02:01 peptide binding pocket. Native MS confirms Leu-2 in the B pocket and Val-9 in the F pocket respectively as the main anchor positions of the pMHC. The analysis of truncated NV9 variants reveals that Val-2 is also a favored binding site. Moreover, Met-5 or Val-5 as well as Val-6 also contribute significantly to binding under certain circumstances.

**Figure 9:**
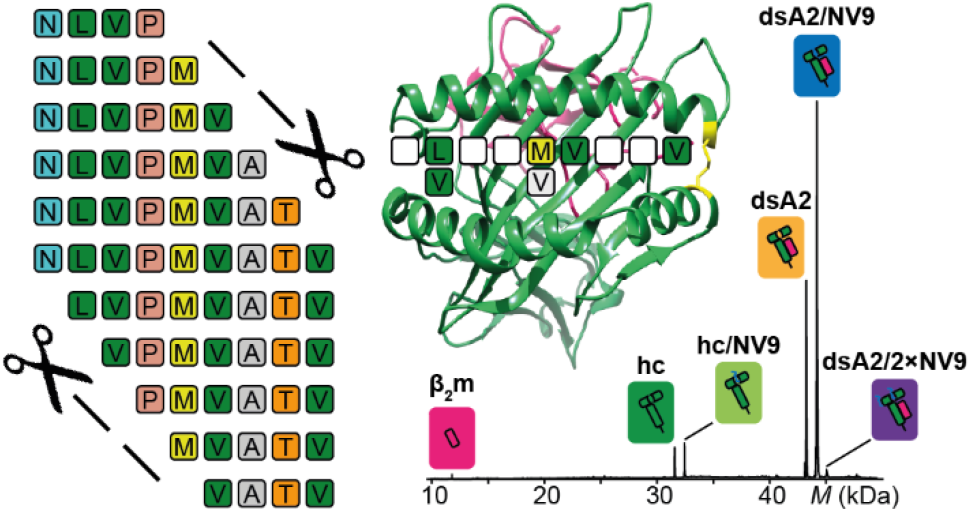
For Table of Contents Only.

Our findings regarding the modification of the peptide termini support the observations of Bouvier *et al*.^7^ that the single-modified nonapeptides can indeed stabilize MHC class I, albeit significantly worse than the non-modified one. However, while no stabilization by the double-modified peptide could be measured there, since no complex was established at all, we succeed in scoring the fully charge-reduced peptide among the other candidates in terms of thermal stabilization and binding strength. Certainly, the comparison of exclusively methylated termini with the acetylation and amidation used in our work is not completely congruent in terms of elimination of hydrogen bonds and occurrence of steric clash. However, acetylation, an intracellular protein degradation signal^51–52^, and amidation, providing the same functional group as an additional amino acid, provide more realistic and biologically relevant modifications. Since *N*-terminal acetylation is one of the most prevalent modifications in pro- and eukaryotes^51, 53^, the observed occupancy for Ac-NV9 in the native MS experiment, which is even higher than for the non-modified NV9 might even be a real effect. Nevertheless, this work confirms that one of the termini needs to be intact to form strong binding to either of the outer pockets. In turn, *N*-terminal peptides from acetylated proteins will likely not bind as decapeptides with a *C*-terminal overhang. Moreover, this explains why undecapeptides with putative overhang on both sides have reduced affinity, as neither of the termini is intact. According to our data, the short, truncated peptides, which are unable to bind using both termini due to spatial limitations, can also stabilize their respective binding pocket individually, which once again supports our thesis concerning the independent stabilization of A and F pocket.

Binding of lipophilic small molecules into the class I binding groove has been shown several times, sometimes quite tightly and (because of alteration of the bound peptidome) with medical consequences^54^. In our MHC class I samples, erucamide is identified as the most prevalent contaminant, most likely from plastic tubes or tips^25^. It appears to bind specifically to the peptide binding groove, since it is displaced by peptides. An alternative explanation would be displacement by a peptide-induced conformational change but is considered less likely, since the empty and the peptide-bound dsA2 are structurally very similar^14^. Thus, erucamide is used as a reference species to estimate the *K*_d_ of the peptides: an increase of dsA2/pep with a simultaneous decrease of dsA2/erucamide demonstrates that the corresponding peptide actually occupies the space in the peptide pocket due to its affinity to dsA2 instead of just clustering non-specifically on the protein. Interestingly, other lipids have been shown to bind class I^55–56^, and since erucamide and related compounds are present in nutritional plants^57^ and produced by *E.coli*^58^, its binding to class I might have biological relevance worth investigating.

HLA-A*02:01 is one of the most prevalent A2 allotypes among Caucasians and Asians, and it has been shown that high binding affinity to A*02:01 correlates with binding affinity to the entire A2 supertype, making it a suitable candidate for comprehensive peptide screening^50^. Epitope-based peptide vaccines offer various advantages with regard to production, stability and mutation risk. However, since HLA is characterized by a strong polymorphism, the search for relevant, allele-spanning peptides has so far been very challenging^50^. Using native MS, we have wide latitude in our approach to identify and analyze potential epitopes. Based on measurements at low acceleration voltages, it is feasible to provide good estimates for the peptides’ binding affinities. Prediction tools such as NetMHC do not provide an option to predict the *K*_d_ for shorter or modified peptides. Furthermore, inconsistencies exist between our data and the affinities predicted using NetMHC. Of course, this is reasonable in terms of absolute values. A smaller share of the observed discrepancy between predicted and approximated dissociation constants can be attributed to an incorrect folding of the protein construct or slight vulnerability against the ammonium acetate solution. Yet, our work shows that the accuracy of *K*_d_ predictions by artificial neural networks needs to be debated. Furthermore, the relative affinities for our fairly large number of peptides are also incongruent with the predictions. Moreover, the affinities determined by native MS and iDSF are in the same order of magnitude and correlate, clearly highlighting issues with the current prediction tool.

Crucially, our measurements at higher energies allow particularly simple evaluation whether a peptide binds strongly or not. The results suggest that pMHC gas-phase stability is mainly determined by side chain interactions within the B and F pockets as well as by binding of the peptide termini. At an acceleration voltage of 50 V, only strongly bound ligands, i.e., those of true biological relevance, are retained. This value hence serves as a cut-off value in our native MS approach that could easily be employed in a native MS based screening. Apart from some attempts^59–61^, to date there is no pervasive high-throughput method for the identification of MHC class I binding peptides with immunogenic potential that is elaborated and reliable. However, such high-throughput screenings are the ultimate key to the development of synthetic peptide vaccines, which in turn offer decisive advantages over conventional vaccines, as there is lesser risk of unwanted host responses, no possibility of reversion to pathogenic phenotypes and no limitation for target diseases^1, 62–63^. Vaccine production is rather easy and detached from the natural source itself, which may be challenging to culture^1, 63^. The greatest advantage of the peptide vaccines lies in their stability. Especially in the context of the SARS-CoV-2 pandemic, mRNA vaccines have become very popular. However, especially these and other conventional vaccines are highly dependent on a continuous cold chain, whereas peptides show long-term thermal stability^1, 62, 64^. This feature is of particular importance, as the need for vaccination is often especially high in tropical and hot climates with a limited medical infrastructure. Nevertheless, mRNA vaccines could also be designed to encode a multitude of MHC-specific peptides. Our work not only provides a valid, sensitive, and rapid method to determine the *K*_d_ of the MHC class I peptide complex but also the basis to develop a novel high-throughput peptide screening method for predicting MHC class I epitopes. Since our technique is based on mass spectrometry, it allows working with very low sample consumption, and it also offers the possibility of simultaneous multi-species analysis.

## MATERIALS AND METHODS

### Production of dsA2 molecules

Production followed Anjanappa, Garcia-Alai *et al*.^14^. HLA-A*02:01(Y84C/A139C) disulfide mutant (dsA2) heavy chain and human β_2_m light chain were expressed in *Escherichia coli* using a pET3a plasmid. The proteins were extracted from inclusion bodies. The dsA2 complex was refolded in presence of 10 mM GM (*Bachem*), concentrated and purified by size-exclusion chromatography on an ÄKTA system (*Cytiva*) using a *Hi-Load 26/600 Superdex 200 pg column* (*Cytiva*).

### Native mass spectrometry

Prior to native MS measurements, *Micro Bio-Spin 6 Columns* (molecular weight cutoff 6 kDa; *Bio-Rad*) were used at 1,000 × g and 4 °C to exchange purified protein samples to 250 mM ammonium acetate (99.99 % purity; *Sigma-Aldrich*), pH 8.0 as buffer surrogate. For native MS experiments, the final concentration of the dsA2 protein was 10 μM. Total peptide concentration (*Genecust*) ranged between 50 μM and 200 μM. No peptidic contaminants either in the bound state or in the free form in the low *m/z* region were detected. Native MS analysis was implemented on a *Q-Tof II* mass spectrometer (*Waters/Micromass*) in positive ESI mode. The instrument was modified to enable high mass experiments (*MS Vision*)^65^. Sample ions were introduced into the vacuum using homemade capillaries via a nano-electrospray ionization source (source pressure: 10 mbar). Borosilicate glass tubes (inner diameter: 0.68 mm, outer diameter: 1.2 mm; *World Precision Instruments*) were pulled into closed capillaries in a two-step program using a squared box filament (2.5 mm × 2.5 mm) within a micropipette puller (*P1000*, *Sutter Instruments*). The capillaries were then gold-coated using a sputter coater (5.0 × 10^-2^ mbar, 30.0 mA, 100 s, 3 runs to vacuum limit 3.0 × 10^-2^ mbar argon, distance of plate holder: 5 cm; *CCU-010*, *safematic*). Capillaries were opened directly on the sample cone of the mass spectrometer. In regular MS mode, spectra were recorded at a capillary voltage of 1.45 kV and a cone voltage of 150 V. Protein species with quaternary structure were assigned by MS/MS analysis. These experiments were carried out using argon as collision gas (1.2 × 10^-2^ mbar). The acceleration voltage ranged from 10 V to 100 V. Comparability of results was ensured as MS quadrupole profiles and pusher settings were kept constant in all measurements. The instrument settings of the mass spectrometer were optimized for non-denaturing conditions. A spectrum of cesium iodide (25 g/L) was recorded on the same day of the particular measurement to calibrate the data.

All spectra were evaluated regarding experimental mass (*MassLynx V4.1*, *Waters*), full width at half maximum (FWHM; *mMass*, Martin Strohalm^66^) and area under the curve (AUC; *UniDec*, Michael T. Marty^67^) of the detected mass species. The values of the shown averaged masses, FWHM (**Table S1**) and AUC (**Table S2**) of the different species as well as the corresponding standard deviation result from at least three independent measurements. Narrow peak widths indicate rather homogeneous samples. In order to eliminate non-specific ESI clustering within the results, the raw data were corrected using the dsA2/pep fraction of the negative control YF9. Affinity *K*_d,high_ was calculated directly from AUC measured at 150 V cone voltage and 10 V acceleration voltage, whereas *K*_d,low_ was indirectly derived from the AUC of the dsA2/erucamide fraction. Since the cone voltage is linearly proportional to the ISD, the distribution of peptide-bound and peptide-unbound dsA2 at non-dissociating conditions can be estimated. According to the equation (*occupancy* = - 0.0048 × *cone voltage* + 1.1747) derived from the linear relationship of cone voltage and occupancy, the binding groove is fully occupied at 36 V and thus reflects the in-solution conditions of the superstoichiometric mixture. The peptide-free (erucamide-bound) fraction of the protein measured at cone voltage of 150 V can be corrected using the linear equation. Subsequently, if this fraction is subtracted from the possible 100 % occupancy, the fraction that binds the peptide is obtained.

### Differential scanning fluorimetry

Thermal stability and binding affinity were determined using nanoscale differential scanning fluorimetry on a *Prometheus NT.48* (*NanoTemper Technologies*). Capillaries were filled with 10 μL of respective samples in duplicates and loaded into the reading chamber. The scan rate was 1 °C/min ranging from 20 °C to 80 °C for thermal stability and 95 °C for binding affinity measurements. Protein unfolding was measured by detecting the temperature-dependent change in intrinsic tryptophan fluorescence at emission wavelengths of 330 nm and 350 nm.

### Thermal stability (nDSF)

2 μM of empty dsA2 were dissolved in citrate phosphate buffer, pH 7.6^68^ and incubated with peptidic ligands (0.2 μM to 2 mM) on ice for 30 min. Melting curves and *T_m_* values were generated by *PR.ThermControl V2.1* software (*NanoTemper Technologies*) using the first derivative of the fluorescence at 330 nm.

### Binding affinity (iDSF)

Empty dsA2 dissolved in 20 mM Tris pH 8, 150 mM NaCl was incubated with peptidic ligands at different concentrations depending on their predicted or assumed *K*_d_ range. For each peptide, a two-fold serial dilution series (11 concentrations) was prepared, while protein concentration was kept constant at 2.2 μM. A pure protein was analyzed as well. *PR.ThermControl V2.1.2* software (*NanoTemper Technologies*) was used to control the device. Data processing and evaluation was executed via the *FoldAffinity* web server (*EMBL Hamburg*^39^).

## Supporting information

complete supplement

## ASSOCIATED CONTENT

### Supporting Information

The following Supporting Information is available free of charge at the ACS website.

**Figure S1.** Small molecule tandem MS analysis verifies that erucamide accounts for a large proportion of the contaminants found in dsA2 samples.

**Figure S2.** Raw spectrum of dsA2/NV9.

**Figure S3.** Melting temperature (*T_m_*) of dsA2 in presence of two corresponding truncated NV9 variants.

**Figure S4.** Overall area under the curve (AUC) for the detected dsA2 mass species at 25 V acceleration voltage.

**Figure S5.** Overall area under the curve (AUC) for the detected dsA2 mass species at 50 V acceleration voltage.

**Table S1.** Experimental masses and FWHM for dsA2 and different peptides obtained by native mass spectrometry.

**Table S2.** Overall area under the curve (AUC) for the detected dsA2 mass species at different acceleration voltages.

**Table S3.** Apparent dissociation constants (*K*_d_) for dsA2 and different peptides obtained by native MS and iDSF.

**Table S4.** Melting temperatures (*T_m_*) for dsA2 and different peptides obtained by nDSF.

The mass spectrometry proteomics data have been deposited to the ProteomeXchange Consortium via the PRIDE^69^ partner repository with the dataset identifier PXD027725.

## Author Contributions

The manuscript was written through contributions of all authors. It was conceptualized by CU, J-DK and SS with methodological advise given by MG-A. CU supervised the project. Experiments were performed by AS, CG, J-DK. Data were analyzed by J-DK and SN, partially using a software provided by SN. Data visualization was conducted by J-DK. The original draft was written by J-DK. Review and editing was done by CU, J-DK and SS with input of all other authors. CU, SS and MG-A acquired funding for the project.

## Notes

Authors declare that they have no competing interests.

## ACKNOWLEDGMENT

The Leibniz Institute for Experimental Virology (HPI) is supported by the Freie und Hansestadt Hamburg and the Bundesministerium für Gesundheit (BMG). J-DK and CU acknowledge support by the Leibniz ScienceCampus InterACt. We thank Raghavendra Anjanappa for the initial protein production and Thomas Dülcks, University of Bremen, for identifying the erucamide in our samples. We acknowledge technical support by the SPC facility at EMBL Hamburg. AS and SS thank Uschi Wellbrock for excellent technical assistance. CU acknowledges the Leibniz Association grant SAW-2014-HPI-4 and SS the Deutsche Forschungsgemeinschaft grant SP583/12-1. Molecular graphics images were produced using the UCSF Chimera package from the Resource for Biocomputing, Visualization, and Informatics at the University of California, San Francisco (supported by NIH P41 RR-01081).

## ABBREVIATIONS

AUC: area under the curve
β_2_m: beta-2 microglobulin
CID: collision-induced dissociation
dsA2: disulfide-stabilized HLA-A*02:01
dsMHC: disulfide-stabilized MHC class I molecules
ESI: electrospray ionization
hc: heavy chain
iDSF: isothermal analysis of nanoscale differential scanning fluorimetry
ISD: in-source dissociation
*K*_d,iDSF_: dissociation constant determined by iDSF
*K*_d,low_ & *K*_d,high_: dissociation constant determined by native MS
*K*_d,th_: dissociation constant predicted by NetMHC
MHC: major histocompatibility complex
MS: mass spectrometry
nDSF: nanoscale differential scanning fluorimetry
PDB: protein Data Bank
pMHC: peptide/MHC class I complex
SEC: size-exclusion chromatography.

