## Supplementary material for "Mapping the peptide binding groove of MHC class I": complete supplement

THIS PDF FILE INCLUDES:

Figures S1 to S5

Tables S1 to S4

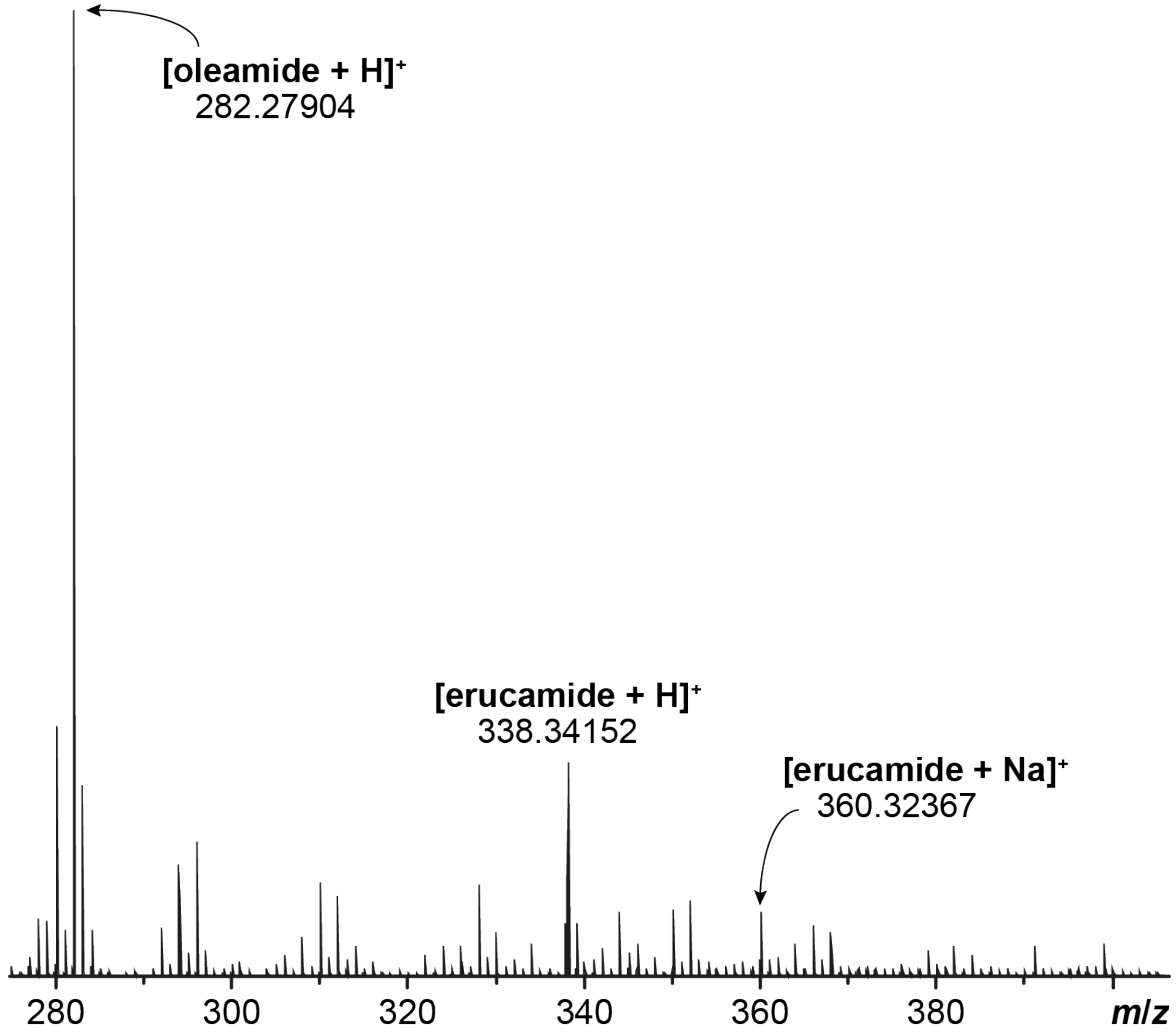


**Figure S1.** Small molecule tandem MS analysis verifies that erucamide accounts for a large proportion of the contaminants found in dsA2 samples. Displayed is a spectrum resulting from the subtraction of the spectrum of plastic ware contaminants (empty vial) from the one of the dsA2 sample itself. Erucamide (337 Da) is hereby identified as the contaminant adducted to dsA2. Although oleamide is also found in the sample, there was no evidence of binding to the protein in the native MS analysis, unlike for erucamide.

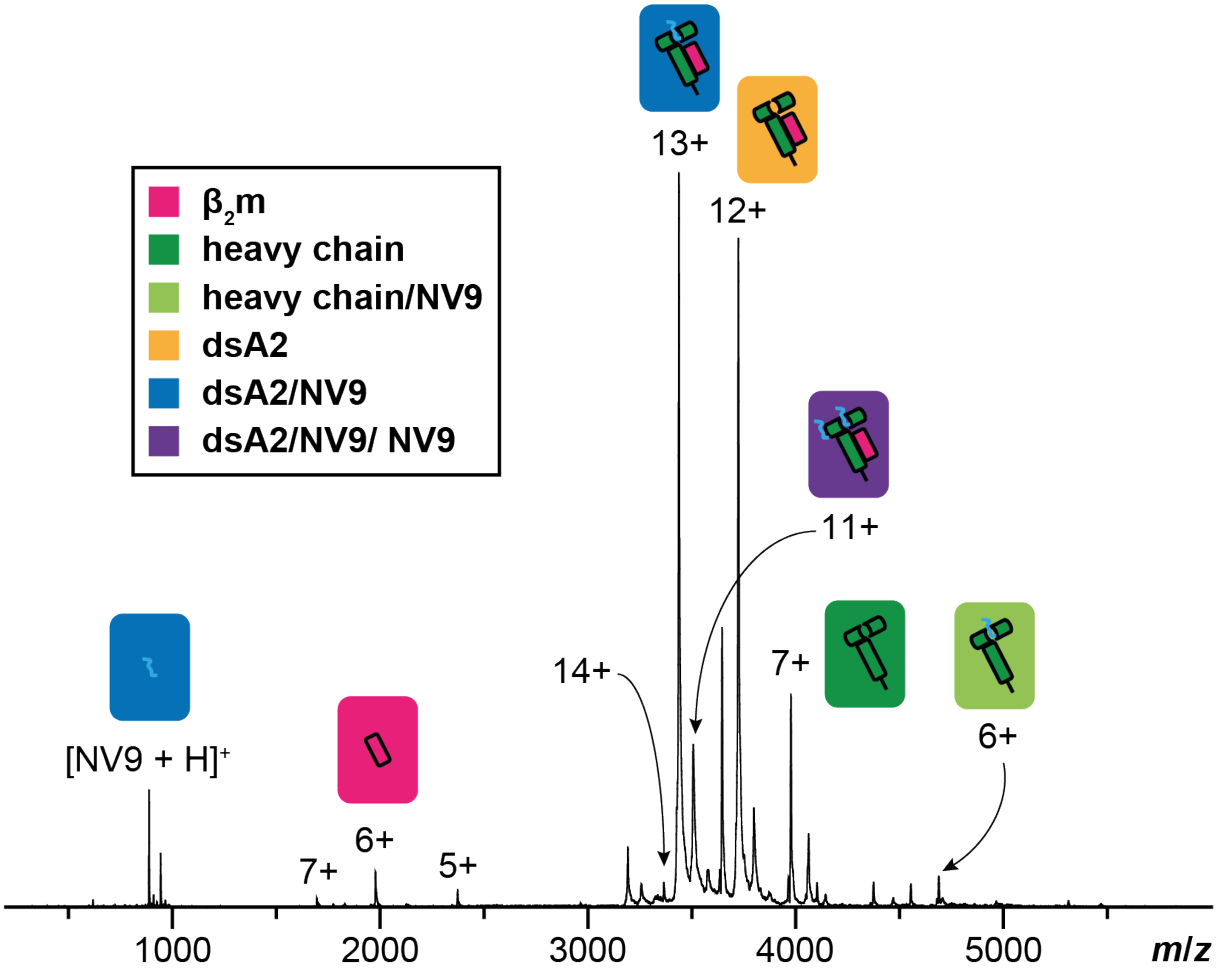


**Figure S2.** Raw spectrum of dsA2/NV9. A native mass spectrum dsA2 in presence of NV9 (protein-peptide ratio 1:5) recorded at an acceleration voltage of 25 V is shown. The dsA2/NV9 is the predominant species (blue). In addition, peptide-free dsA2 (yellow), dissociated β2m (pink) and heavy chain (green) as well as free NV9 can be seen.

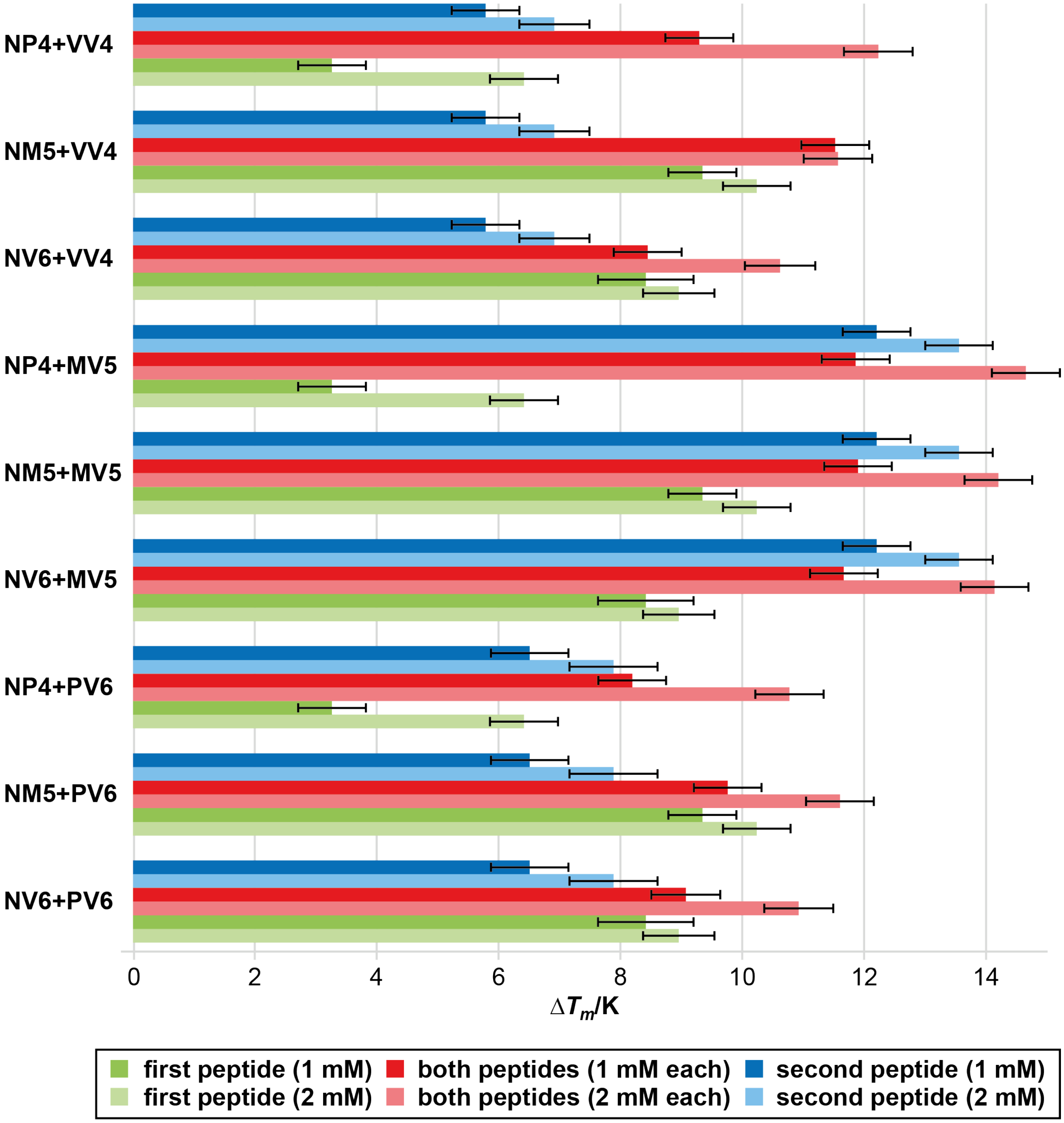


**Figure S3.** Melting temperature ($T_m$) of dsA2 in presence of two corresponding truncated NV9 variants. Nanoscale differential scanning fluorimetry is employed to study thermal denaturation of dsA2 in presence of two peptides at once. 2 µM dsA2 are combined with either exclusively the *N*- (green) or *C*-terminal peptide (blue) or both peptides together (red) at different concentrations. Results for $\Delta T_m$ along with respective standard deviations are displayed.

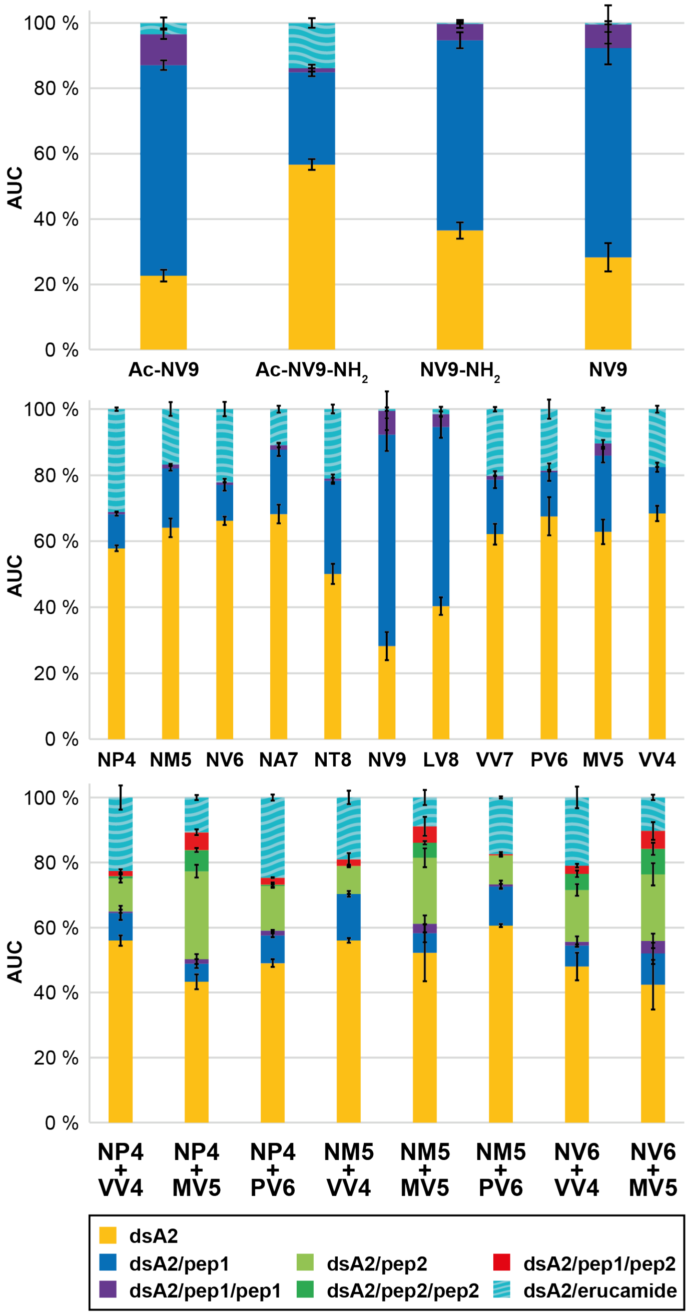


**Figure S4.** Overall area under the curve (AUC) for the detected dsA2 mass species at 25 V acceleration voltage. The AUC is determined over the entire spectrum for the respective mass species at 25 V. The mean value of the AUC in absence or presence of the different peptides (protein-peptide ratio 1:5, 1:10:10 in dual peptide approach) from at least three independent measurements is depicted along with error bars that represent the corresponding standard deviation. "dsA2" (yellow bars) corresponds to the empty HLA-A*02:01(Y84C/A139C) disulfide mutant complex, "dsA2/pep" (blue bars) to dsA2 bound to one peptide, "dsA2/pep/pep" to dsA2 bound to two molecules of this certain peptide (purple bars), "dsA2/pep2" to dsA2 bound to another peptide when two different peptides where present (light green), "dsA2/pep2/pep2" to dsA2 bound to two molecules of the second peptide (dark green), "dsA2/pep1/pep2" to dsA2 bound to one molecule of each of both peptides (red bars) and "dsA2/erucamide" to dsA2 bound to the erucamide (turquoise-striped bars) respectively.

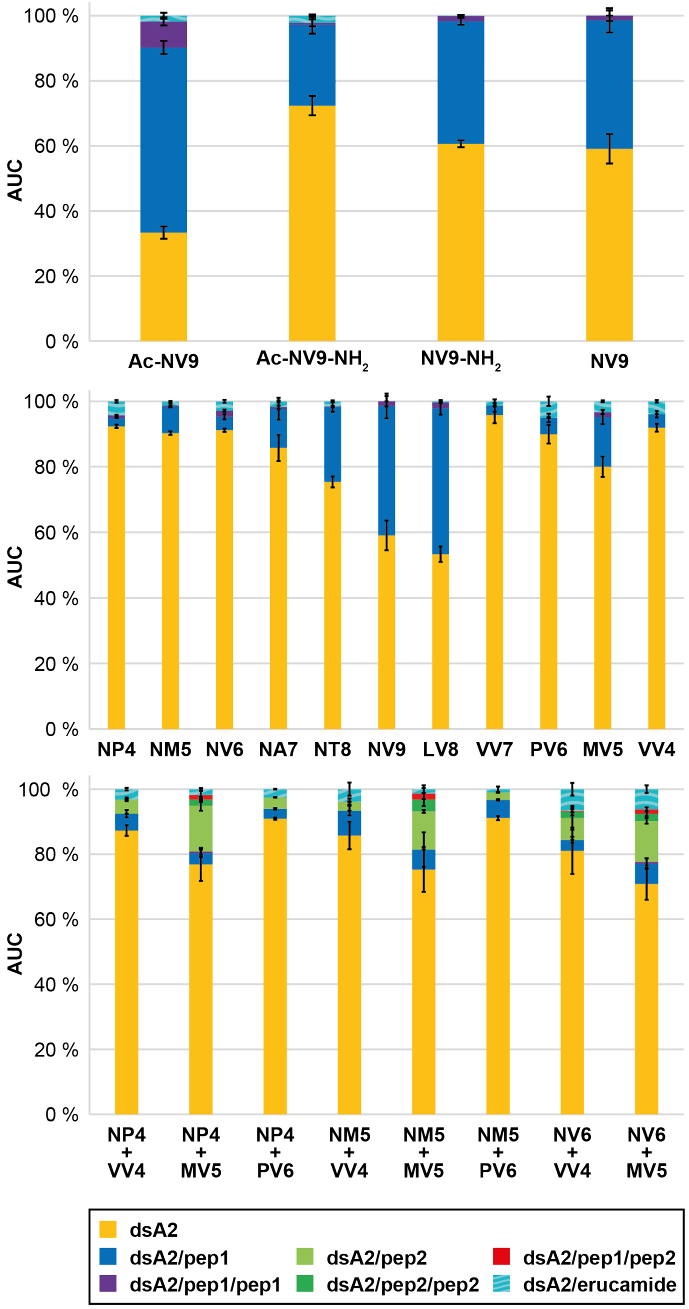


**Figure S5.** Overall area under the curve (AUC) for the detected dsA2 mass species at 50 V acceleration voltage. The AUC is determined over the entire spectrum for the respective mass species 50 V. The mean value of the AUC in absence or presence of the different peptides (protein-peptide ratio 1:5, 1:10:10 in dual peptide approach) from at least three independent measurements is depicted along with error bars that represent the corresponding standard deviation. “dsA2” (yellow bars) corresponds to the empty HLA-A*02:01(Y84C/A139C) disulfide mutant complex, “dsA2/pep” (blue bars) to dsA2 bound to one peptide, “dsA2/pep/pep” to dsA2 bound to two molecules of this certain peptide (purple bars), “dsA2/pep2” to dsA2 bound to another peptide when two different peptides where present (light green), “dsA2/pep2/pep2” to dsA2 bound to two molecules of the second peptide (dark green), “dsA2/pep1/pep2” to dsA2 bound to one molecule of each of both peptides (red bars) and “dsA2/erucamide” to dsA2 bound to the erucamide (turquoise-striped bars) respectively.

**Table S1.** Experimental masses and FWHM for dsA2 and different peptides obtained by native mass spectrometry. Experimental masses ($m_{exp}$) of the different protein species of disulfide stabilized HLA-A*02:01(Y84C/A139C) disulfide mutant (dsA2) in absence or presence of the different peptides (protein-peptide ratio 1:5, 1:10:10 in dual peptide approach) are determined from at least three independent mass spectrometry measurements. They are listed together with the respective values for standard deviation *s* and average full width of the peak at half maximum (FWHM) along with the theoretically calculated molecular weight (*M*). FWHM values are given for the whole peak area where individual species are not fully resolved. [1]NV9 – high affinity control, [2]YF9 – low affinity control, [3]GV9 – minimal binding motif, [4]Ac-NV9 – modified *N*-terminus, [5]Ac-NV9 $NH_2$ – modified *N*- and *C*-terminus, [6]NV9-$NH_2$ – modified *C*-terminus.

| mass species | $M$/Da | $m_{exp}$/Da | $s(m_{exp})$/Da | FWHM/Da | $s(m_{exp})$/Da |
|---|---|---|---|---|---|
| dsA2 - M1 | 43,608 | 43,603 | 4 | 20 | 20 |
| dsA2 | 43,739 | 43,733 | 4 | 3 | 2 |
| heavy chain (hc) | 31,877 | 31,873 | 2 | 2.1 | 0.3 |
| $\beta_2$m chain - M1 | 11,731 | 11,729 | 2 | 1.1 | 0.4 |
| $\beta_2$m chain | 11,862 | 11,860 | 1 | 1.4 | 0.2 |
| dsA2/NV9[1] | 44,682 | 44,678 | 1 | 3 | 2 |
| dsA2/2×NV9 | 45,625 | 45,624 | 4 | 12 | 6 |
| hc/NV9 | 32,820 | 32,816.9 | 0.8 | 2.2 | 0.3 |
| dsA2/YF9[2] | 44,856 | 44,849 | 1 | 7 | 3 |
| dsA2/GV9[3] | 44,369 | 44,363.8 | 0.7 | 2 | 1 |
| dsA2/2×GV9 | 44,999 | 44,995 | 2 | 3 | 2 |
| hc/GV9 | 32,507 | 32,503.0 | 0.5 | 2.08 | 0.09 |
| dsA2/Ac-NV9[4] | 44,724 | 44,718.3 | 0.5 | 1.6 | 0.3 |
| dsA2/2×Ac-NV9 | 45,709 | 45,704 | 2 | 3 | 2 |
| hc/Ac-NV9 | 32,862 | 32,857.3 | 0.5 | 1.9 | 0.1 |
| dsA2/Ac-NV9-$NH_2$[5] | 44,723 | 44,717.4 | 1.0 | 1.4 | 0.1 |
| dsA2/2×Ac-NV9-$NH_2$ | 45,707 | 45,701.00 | 1.00 | 1.8 | 0.6 |
| hc/Ac-NV9-$NH_2$ | 32,861 | 32,857.0 | 0.9 | 1.7 | 0.2 |
| dsA2/NV9-$NH_2$[6] | 44,681 | 44,676.4 | 0.5 | 1.38 | 0.06 |
| dsA2/2×NV9-$NH_2$ | 45,623 | 45,619.6 | 0.9 | 1.6 | 0.6 |
| hc/NV9-$NH_2$ | 32,819 | 32,815.1 | 0.9 | 1.72 | 0.10 |

| mass species | $M$/Da | $m_{exp}$/Da | $s(m_{exp})$/Da | FWHM/Da | $s(m_{exp})$/Da |
|---|---|---|---|---|---|
| dsA2/NP4 | 44,181 | 44,176 | 3 | 10 | 10 |
| dsA2/2×NP4 | 44,623 | 44,619 | 4 | 30 | 40 |
| hc/NP4 | 32,319 | 32,317 | 2 | 20 | 40 |
| dsA2/NM5 | 44,312 | 44,309 | 2 | 6 | 3 |
| dsA2/2×NM5 | 44,885 | 44,883 | 2 | 60 | 50 |
| hc/NM5 | 32,450 | 32,449 | 1 | 2.5 | 0.6 |
| dsA2/NV6 | 44,411 | 44411 | 1 | 7 | 2 |
| dsA2/2×NV6 | 45,083 | 45,082 | 3 | 145 | 6 |
| hc/NV6 | 32,549 | 32,547 | 3 | 6 | 7 |
| dsA2/NA7 | 44,481 | 44,476.4 | 0.5 | 5 | 3 |
| dsA2/2×NA7 | 45,223 | 45,225 | 4 | 30 | 60 |
| hc/NA7 | 32,619 | 32,615 | 2 | 2.4 | 0.5 |
| dsA2/NT8 | 44,583 | 44,579.33 | 1.00 | 3 | 2 |
| dsA2/2×NT8 | 45,427 | 45,425 | 2 | 7 | 5 |
| hc/NT8 | 32,721 | 32,718.1 | 0.3 | 2.4 | 0.7 |
| dsA2/LV8 | 44,568 | 44,565.8 | 0.7 | 2.6 | 0.7 |
| dsA2/2×LV8 | 45,397 | 45,396.1 | 0.3 | 3.4 | 1.0 |
| hc/LV8 | 32,706 | 32,704.8 | 0.4 | 2.18 | 0.04 |
| dsA2/VV7 | 44,455 | 44,447.8 | 0.8 | 7 | 3 |
| dsA2/2×VV7 | 45,171 | 45,166 | 3 | 3 | 1 |
| hc/VV7 | 32,593 | 32,586 | 3 | 3 | 2 |
| dsA2/PV6 | 44,356 | 44,350 | 3 | 5 | 3 |
| dsA2/2×PV6 | 44,973 | 44969 | 4 | 100 | 50 |
| hc/PV6 | 32,194 | 32,487 | 3 | 2.3 | 0.4 |
| dsA2/MV5 | 44,259 | 44,254 | 3 | 8 | 12 |
| dsA2/2×MV5 | 44,779 | 44,774 | 4 | 20 | 30 |

| mass species | $M$/Da | $m_{exp}$/Da | $s(m_{exp})$/Da | FWHM/Da | $s(m_{exp})$/Da |
|---|---|---|---|---|---|
| hc/MV5 | 32,397 | 32,388.4 | 1.0 | 2.1 | 0.2 |
| dsA2/VV4 | 44,127 | 44,125 | 5 | 10 | 10 |
| dsA2/2×VV4 | 44,515 | 44,512 | 8 | 20 | 30 |
| hc/VV4 | 32,265 | 32,255 | 1 | 10 | 3 |
| dsA2/erucamide | 44,077 | 44,071 | 5 | 10 | 20 |
| dsA2/NP4/VV4 | 44,569 | 44,565 | 2 | 30 | 40 |
| dsA2/NP4/MV5 | 44,701 | 44,697 | 2 | 4 | 5 |
| dsA2/NP4/PV6 | 44,798 | 44,792 | 3 | 7 | 6 |
| dsA2/NM5/VV4 | 44,700 | 44,699 | 2 | 30 | 40 |
| dsA2/NM5/MV5 | 44,832 | 44,827 | 2 | 12 | 3 |
| dsA2/NM5/PV6 | 44,929 | 44,927 | 2 | 100 | 40 |
| dsA2/NV6/VV4 | 44,799 | 44,797 | 4 | 30 | 40 |
| dsA2/NV6/MV5 | 45,451 | 44927 | 5 | 100 | 60 |

**Table S2.** Overall area under the curve (AUC) for the detected dsA2 mass species at different acceleration voltages. The AUC is determined over the entire spectrum for the respective mass species at 10 V, 25 V and 50 V. The mean value of the AUC in absence or presence of the different peptides (protein-peptide ratio 1:5, 1:10:10 in dual peptide approach) from at least three independent measurements is listed together with their corresponding standard deviation *s*. "dsA2" corresponds to the empty HLA-A*02:01(Y84C/A139C) disulfide mutant complex, "dsA2/pep" to dsA2 bound to one peptide, "dsA2/pep/pep" to dsA2 bound to two molecules of this certain peptide, "dsA2/pep2" to dsA2 bound to another peptide when two different peptides where present, "dsA2/pep2/pep2" to dsA2 bound to two molecules of the second peptide, "dsA2/pep1/pep2" to dsA2 bound to one molecule of each of both peptides and "dsA2/erucamide" to dsA2 bound to the erucamide respectively. [1]NV9 – high affinity control, [2]YF9 – low affinity control, [3]GV9 – minimal binding motif, [4]Ac-NV9 – modified *N*-terminus, [5]Ac-NV9 $NH_2$ – modified *N*- and *C*-terminus, [6]NV9-$NH_2$ – modified *C*-terminus.

| peptide | acc. volt. | dsA2 | | dsA2/pep | | dsA2/pep/pep | | dsA2/pep2 | | dsA2/pep2/pep2 | | dsA2/pep1/pep2 | | dsA2/erucamide | |
|---|---|---|---|---|---|---|---|---|---|---|---|---|---|---|---|
| | | AUC | s | AUC | s | AUC | s | AUC | s | AUC | s | AUC | s | AUC | s |
| NV9[1] | 10 V | 31% | 2% | 64% | 3% | 4% | 4% | | | | | | | 2% | 2% |
| | 25 V | 28% | 4% | 64% | 5% | 7% | 6% | | | | | | | 0.5% | 0.5% |
| | 50 V | 59% | 5% | 40% | 4% | 1% | 2% | | | | | | | 0% | 0% |
| YF9[2] | 10 V | 56% | 3% | 4% | 2% | 0% | 0% | | | | | | | 39% | 2% |
| | 25 V | 51% | 4% | 5% | 2% | 0% | 0% | | | | | | | 44% | 3% |
| | 50 V | 95% | 5% | 5% | 2% | 0% | 0% | | | | | | | 2% | 2% |
| GV9[3] | 10 V | 52% | 2% | 43% | 2% | 1.5% | 0.4% | | | | | | | 4.3% | 0.6% |
| | 25 V | 50% | 1% | 43% | 3% | 2.2% | 0.9% | | | | | | | 5% | 2% |
| | 50 V | 62% | 3% | 35% | 3% | 1.1% | 0.3% | | | | | | | 1.7% | 0.2% |
| Ac-NV9[4] | 10 V | 24% | 2% | 63% | 3% | 11% | 2% | | | | | | | 2.1% | 0.5% |
| | 25 V | 23% | 2% | 64% | 1% | 9% | 1% | | | | | | | 3% | 2% |
| | 50 V | 33% | 2% | 57% | 2% | 8% | 1% | | | | | | | 1.7% | 0.9% |
| Ac-NV9-NH2[5] | 10 V | 55% | 2% | 27.9% | 0.5% | 2.4% | 0.6% | | | | | | | 15% | 1% |
| | 25 V | 57% | 2% | 28% | 1% | 1% | 1% | | | | | | | 14% | 1% |
| | 50 V | 72% | 3% | 25% | 3% | 1% | 1% | | | | | | | 2.2% | 0.4% |
| NV9-NH2[6] | 10 V | 36% | 2% | 59% | 2% | 5% | 1% | | | | | | | 0.4% | 0.2% |
| | 25 V | 36% | 2% | 58% | 2% | 5% | 1% | | | | | | | 0.3% | 0.3% |
| | 50 V | 61% | 1% | 38% | 1% | 1.6% | 0.3% | | | | | | | 0.1% | 0.2% |

| peptide | acc. volt. | dsA2 | | dsA2/pep | | dsA2/pep/pep | | dsA2/pep2 | | dsA2/pep2/pep2 | | dsA2/pep1/pep2 | | dsA2/erucamide | |
|---|---|---|---|---|---|---|---|---|---|---|---|---|---|---|---|
| | | AUC | s | AUC | s | AUC | s | AUC | s | AUC | s | AUC | s | AUC | s |
| NP4 | 10 V | 55% | 2% | 11% | 1% | 0.6% | 0.2% | | | | | | | 33.8% | 0.6% |
| | 25 V | 57.8% | 0.9% | 10.4% | 0.5% | 0.69% | 0.06% | | | | | | | 31.1% | 0.5% |
| | 50 V | 92.4% | 0.5% | 2.6% | 0.1% | 0.8% | 0.1% | | | | | | | 4.3% | 0.3% |
| NM5 | 10 V | 62% | 2% | 17.9% | 0.6% | 1.0% | 0.2% | | | | | | | 19% | 1% |
| | 25 V | 64% | 3% | 18.0% | 0.7% | 1.2% | 0.2% | | | | | | | 17% | 2% |
| | 50 V | 90.3% | 0.6% | 8.1% | 0.3% | 0.3% | 0.6% | | | | | | | 1.3% | 0.1% |
| NV6 | 10 V | 66% | 1% | 10% | 2% | 0.5% | 0.1% | | | | | | | 24% | 1% |
| | 25 V | 66% | 1% | 11% | 2% | 0.69% | 0.08% | | | | | | | 22% | 2% |
| | 50 V | 91.2% | 0,5% | 4.1% | 0.8% | 1.9% | 0.3% | | | | | | | 2.9% | 0.4% |
| NA7 | 10 V | 75% | 3% | 16% | 2% | 0.8% | 0.3% | | | | | | | 8% | 1% |
| | 25 V | 68% | 3% | 19% | 2% | 1.4% | 0.7% | | | | | | | 11% | 1% |
| | 50 V | 86% | 4% | 12% | 3% | 0.5% | 0.5% | | | | | | | 1.8% | 0.2% |
| NT8 | 10 V | 47.3% | 0.9% | 28% | 1% | 0.6% | 0.5% | | | | | | | 24% | 1% |
| | 25 V | 50% | 3% | 28.2% | 0.9% | 1% | 1% | | | | | | | 21% | 1% |
| | 50 V | 75% | 2% | 23% | 1% | 0.2% | 0.3% | | | | | | | 1.6% | 0.3% |
| LV8 | 10 V | 42% | 6% | 54% | 5% | 2.8% | 0.3% | | | | | | | 1.7% | 0.9% |
| | 25 V | 40% | 3% | 54% | 3% | 3.9% | 0.8% | | | | | | | 1.5% | 0.7% |
| | 50 V | 53% | 2% | 45% | 2% | 1.6% | 0.2% | | | | | | | 0.4% | 0.4% |
| VV7 | 10 V | 61.8% | 0.5% | 17% | 2% | 1.7% | 0.3% | | | | | | | 19% | 3% |
| | 25 V | 62% | 3% | 17% | 3% | 1.16% | 0.06% | | | | | | | 20.2% | 0.7% |
| | 50 V | 96% | 3% | 3% | 2% | 0% | 0% | | | | | | | 1.3% | 0.6% |
| PV6 | 10 V | 68% | 6% | 12% | 2% | 0.5% | 0.6% | | | | | | | 19% | 3% |
| | 25 V | 68% | 6% | 13% | 3% | 0.5% | 0.2% | | | | | | | 19% | 3% |
| | 50 V | 90% | 3% | 5% | 1% | 0.1% | 0.1% | | | | | | | 5% | 1% |

| peptide | acc. volt. | dsA2 | | dsA2/pep | | dsA2/pep/pep | | dsA2/pep2 | | dsA2/pep2/pep2 | | dsA2/pep1/pep2 | | dsA2/erucamide | |
|---|---|---|---|---|---|---|---|---|---|---|---|---|---|---|---|
| | | AUC | s | AUC | s | AUC | s | AUC | s | AUC | s | AUC | s | AUC | s |
| MV5 | 10 V | 64% | 3% | 23% | 2% | 3.2% | 0.7% | | | | | | | 10.4% | 0.2% |
| | 25 V | 63% | 4% | 23% | 2% | 3.7% | 1.0% | | | | | | | 10.4% | 0.4% |
| | 50 V | 80% | 3% | 15% | 2% | 1.5% | 0.6% | | | | | | | 3.3% | 0.3% |
| VV4 | 10 V | 68% | 1% | 14% | 1% | 0% | 0% | | | | | | | 18.093% | 0.007% |
| | 25 V | 68% | 2% | 14% | 1% | 0% | 0% | | | | | | | 17.6% | 1.0% |
| | 50 V | 92% | 1% | 4.1% | 1.0% | 0% | 0% | | | | | | | 4.0% | 0.4% |
| NP4 | 10 V | 54% | 2% | 9% | 3% | 1.1% | 0.5% | 10% | 1% | 1.2% | 1.0% | 2.1% | 0.5% | 22% | 4% |
| + | 25 V | 56% | 2% | 8% | 2% | 0.5% | 0.5% | 10% | 1% | 0.7% | 0.7% | 1.6% | 0.5% | 22% | 4% |
| VV4 | 50 V | 87% | 2% | 5% | 1% | 0% | 0% | 4.4% | 0.6% | 0% | 0% | 0% | 0% | 3.1% | 0.4% |
| NP4 | 10 V | 42.9% | 0.9% | 6% | 2% | 1,8% | 0.3% | 27% | 1% | 5.6% | 0.3% | 5.2% | 0.8% | 10% | 1% |
| + | 25 V | 43% | 2% | 5% | 1% | 1% | 2% | 27% | 2% | 6.4% | 0.6% | 5.3% | 0.9% | 10.5% | 0.8% |
| MV5 | 50 V | 76% | 5% | 3% | 1% | 0.5% | 0.9% | 14% | 2% | 1.9% | 0.8% | 1% | 1% | 1.8% | 0.3% |
| NP4 | 10 V | 48% | 3% | 8% | 1% | 1.7% | 0.3% | 13% | 2% | 0.9% | 0.8% | 2.8% | 0.2% | 25% | 2% |
| + | 25 V | 49% | 1% | 8.5% | 0.6% | 1.4% | 0.2% | 13.6% | 0.5% | 0.4% | 0.7% | 2.1% | 0.1% | 24.5% | 0.9% |
| PV6 | 50 V | 90.8% | 0.3% | 3.0% | 0.3% | 0% | 0% | 3.5% | 0.1% | 0% | 0% | 0% | 0% | 2.5% | 0.1% |
| NM5 | 10 V | 56% | 3% | 13% | 2% | 1% | 1% | 8.9% | 0.8% | 0.5% | 0.8% | 1.8% | 1.6% | 18% | 4% |
| + | 25 V | 55.7% | 0.7% | 14.2% | 0.9% | 0% | 0% | 8.6% | 0.4% | 0% | 0% | 2.0% | 1.9% | 19% | 2% |
| VV4 | 50 V | 86% | 4% | 8% | 1% | 0% | 0% | 2.8% | 1.0% | 0% | 0% | 0% | 0% | 4% | 2% |
| NM5 | 10 V | 46% | 7% | 8% | 3% | 5% | 4% | 20% | 1% | 4.9% | 0.8% | 6% | 1% | 9% | 1% |
| + | 25 V | 51% | 9% | 6% | 3% | 3% | 3% | 20% | 3% | 4.5% | 0.5% | 5% | 3% | 9% | 2% |
| MV5 | 50 V | 75% | 7% | 6% | 5% | 0% | 0% | 11.7% | 0.4% | 4% | 2% | 2% | 2% | 1% | 1% |
| NM5 | 10 V | 58% | 2% | 11.9% | 0.6% | 1% | 2% | 9% | 2% | 0% | 0% | 0.4% | 0.7% | 18% | 1% |
| + | 25 V | 60.3% | 0.5% | 12.0% | 0.8% | 1% | 1% | 8.9% | 0.4% | 0% | 0% | 0.4% | 0.7% | 17.3% | 0.4% |
| PV6 | 50 V | 91.1% | 0.7% | 5.6% | 0.2% | 0% | 0% | 2.4% | 0.2% | 0% | 0% | 0% | 0% | 0.9% | 0.8% |

| peptide | acc. volt. | dsA2 | | dsA2/pep | | dsA2/pep/pep | | dsA2/pep2 | | dsA2/pep2/pep2 | | dsA2/pep1/pep2 | | dsA2/erucamide | |
|---|---|---|---|---|---|---|---|---|---|---|---|---|---|---|---|
| | | AUC | s | AUC | s | AUC | s | AUC | s | AUC | s | AUC | s | AUC | s |
| NV6 | 10 V | 47% | 8% | 6% | 1% | 1% | 0% | 17% | 4% | 7% | 4% | 3% | 0% | 18% | 1% |
| + | 25 V | 48% | 4% | 6% | 0% | 1% | 2% | 16% | 2% | 5% | 1% | 2% | 1% | 21% | 3% |
| VV4 | 50 V | 81% | 7% | 3% | 1% | 0% | 0% | 7% | 4% | 2% | 1% | 0% | 0% | 7% | 2% |
| NV6 | 10 V | 47% | 5% | 8% | 2% | 4% | 2% | 21% | 1% | 6.5% | 0.8% | 3.0% | 0.3% | 10% | 2% |
| + | 25 V | 42% | 8% | 9% | 3% | 4% | 2% | 20% | 3% | 8% | 2% | 5% | 3% | 10.2% | 0.8% |
| MV5 | 50 V | 71% | 5% | 6% | 2% | 1% | 1% | 12.5% | 0.9% | 2.1% | 0.7% | 1.4% | 0.6% | 6% | 1% |

**Table S3.** Apparent dissociation constants ($K_d$) for dsA2 and different peptides obtained by native MS and iDSF. The $K_d$ is calculated from the respective area under the curve values at 10 V acceleration voltage (protein-peptide ratio 1:5). $K_{d,high}$ is an affinity determined on the basis of a real experiment at a cone voltage of 150 V, while a theoretical cone voltage of 36 V is assumed for $K_{d,low}$ to correct for ion-source decay. $K_{d,iDSF}$ is derived by two independent measurements. Protein concentration is 2.2 µM. For each ligand, a two-fold serial dilution series is prepared using 11 concentrations depending on their predicted or assumed $K_d$ range. The listed standard deviation *s* for $K_d$ is determined using common equations, which estimate the propagation of uncertainty. [1]NV9 – high affinity control, [2]GV9 – minimal binding motif, [3]Ac-NV9 – modified *N*-terminus, [4]Ac-NV9 $NH_2$ – modified *N*- and *C*-terminus, [5]NV9-$NH_2$ – modified *C*-terminus; *iDSF reaches its limits at affinities below 200 nm, hence the values of grayed-out peptides are not reliable.

| peptide | sequence | $K_{d,high}$ | | $K_{d,low}$ | | $K_{d,iDSF}$ | |
|---|---|---|---|---|---|---|---|
| | | $K_d$/µM | s/µM | $K_d$/µM | s/µM | $K_d$/µM | s/µM |
| NV9[1] | NLVPMVATV | 8 | 2 | 0.06 | 0.08 | 0.04* | 0.01 |
| GV9[2] | GLGGGGGGV | 35 | 2 | 0.5 | 0.2 | 0.36 | 0.06 |
| Ac-NV9[3] | | 4.5 | 1.0 | 0.11 | 0.05 | 0.61 | 0.08 |
| Ac-NV9-NH2[4] | | 80 | 7 | 7 | 3 | 4 | 1 |
| NV9-NH2[5] | | 10 | 1 | 0.004 | 0.003 | 0.001* | 0.001 |
| NP4 | NLVP | 350 | 60 | 100 | 70 | 50 | 20 |
| NM5 | NLVPM | 180 | 30 | 15 | 6 | 11 | 3 |
| NV6 | NLVPMV | 380 | 70 | 30 | 10 | 9 | 2 |
| NA7 | NLVPMVA | 210 | 40 | 2.0 | 0.7 | 3.6 | 0.5 |
| NT8 | NLVPMVAT | 92 | 5 | 30 | 10 | 2.6 | 0.4 |
| LV8 | LVPMVATV | 17 | 3 | 0.07 | 0.06 | 0.008* | 0.005 |
| VV7 | VPMVATV | 180 | 30 | 15 | 7 | 7.8 | 1.0 |
| PV6 | PMVATV | 300 | 200 | 15 | 7 | 15 | 2 |
| MV5 | MVATV | 110 | 10 | 3 | 1 | 1.6 | 0.2 |
| VV4 | VATV | 270 | 30 | 13 | 4 | 50 | 20 |
| NP4+VV4 | | | | 90 | 30 | | |
| NP4+MV5 | | | | 11 | 3 | | |
| NP4+PV6 | | | | 130 | 50 | | |
| NM5+VV4 | | | | 50 | 20 | | |
| NM5+MV5 | | | | 9 | 3 | | |

| peptide | sequence | $K_{d,high}$ | | $K_{d,low}$ | | $K_{d,iDSF}$ | |
|---|---|---|---|---|---|---|---|
| | | $K_d$/µM | s/µM | $K_d$/µM | s/µM | $K_d$/µM | s/µM |
| NM5+PV6 | | | | 50 | 10 | | |
| NV6+VV4 | | | | 50 | 10 | | |
| NV6+MV5 | | | | 13 | 4 | | |

**Table S4.** Melting temperatures ($T_m$) for dsA2 and different peptides obtained by nDSF. The $T_m$ as well as the resulting *s* for dsA2 in absence or presence of the different peptides is defined by at least two independent measurements. Protein concentration is 2 µM. [1]NV9 – high affinity control, [2]YF9 – low affinity control, [3]GV9 – minimal binding motif, [4]Ac-NV9 – modified *N*-terminus, [5]Ac-NV9 $NH_2$ – modified *N*- and *C*-terminus, [6]NV9-$NH_2$ – modified *C*-terminus.

| **peptide** | **0.2 µM** | | **2 µM** | | **20 µM** | | **1 mM** | | **2 mM** | |
|---|---|---|---|---|---|---|---|---|---|---|
| | **$T_m$/°C** | **s/°C** | **$T_m$/°C** | **s/°C** | **$T_m$/°C** | **s/°C** | **$T_m$/°C** | **s/°C** | **$T_m$/°C** | **s/°C** |
| empty | 35.7 | 0.6 | | | | | | | | |
| NV9[1] | 35.8 | 0.8 | 58.96 | 0.07 | 59.1 | 0.1 | 60.698 | 0.007 | | |
| YF9[2] | 36.1 | 0.8 | 36.1 | 0.8 | 36.2 | 0.8 | | | | |
| GV9[3] | 36.4 | 0.6 | 38.8 | 0.4 | 42.1 | 0.2 | 47.6 | 0.1 | | |
| Ac-NV9[4] | 36.6 | 0.7 | 39.6 | 0.6 | 43.2 | 0.6 | 48.420 | 0.001 | | |
| Ac-NV9-NH2[5] | 36.0 | 0.8 | 36.6 | 0.4 | 38.4 | 0.3 | | | | |
| NV9-NH2[6] | 36.0 | 0.9 | 46.9 | 0.2 | 50.3 | 0.3 | 55.53 | 0.04 | | |
| NP4 | 35.6 | 0.4 | 35.7 | 0.5 | 35.8 | 0.4 | 38.95 | 0.04 | 42.10 | 0.06 |
| NM5 | 35.6 | 0.5 | 35.9 | 0.4 | 37.3 | 0.3 | 45.04 | 0.07 | 45.93 | 0.03 |
| NV6 | 35.5 | 0.1 | 35.9 | 0.2 | 37.61 | 0.05 | 44.1 | 0.6 | 44.6 | 0.2 |
| NA7 | 35.51 | 0.09 | 35.9 | 0.1 | 37.95 | 0.06 | 45.442 | 0.007 | | |
| NT8 | 35.5 | 0.1 | 36.02 | 0.09 | 38.78 | 0.02 | 46.89 | 0.02 | | |
| LV8 | 35.40 | 0.08 | 43.5 | 0.1 | 47.28 | 0.04 | 53.30 | 0.04 | | |
| VV7 | 35.5 | 0.1 | 35.7 | 0.2 | 37.0 | 0.1 | 42.29 | 0.01 | | |
| PV6 | 35.5 | 0.2 | 35.62 | 0.09 | 36.40 | 0.02 | 42.2 | 0.3 | 43.6 | 0.5 |
| MV5 | 35.64 | 0.08 | 36.5 | 0.1 | 39.31 | 0.07 | 47.90 | 0.01 | 49.253 | 0.003 |
| VV4 | 35.475 | 0.002 | 35.1 | 0.6 | 35.5 | 0.2 | 41.47 | 0.03 | 42.6 | 0.1 |
| NP4+VV4 | | | | | | | 44.99 | 0.05 | 47.93 | 0.10 |
| NP4+MV5 | | | | | | | 47.55 | 0.06 | 50.35 | 0.05 |
| NP4+PV6 | | | | | | | 43.88 | 0.05 | 46.47 | 0.08 |
| NM5+VV4 | | | | | | | 47.22 | 0.05 | 47.26 | 0.08 |
| NM5+MV5 | | | | | | | 47.59 | 0.03 | 49.900 | 0.002 |
| NM5+PV6 | | | | | | | 45.45 | 0.02 | 47.295 | 0.010 |

| peptide | 0.2 µM | | 2 µM | | 20 µM | | 1 mM | | 2 mM | |
|---|---|---|---|---|---|---|---|---|---|---|
| | $T_m$/°C | s/°C | $T_m$/°C | s/°C | $T_m$/°C | s/°C | $T_m$/°C | s/°C | $T_m$/°C | s/°C |
| NV6+VV4 | | | | | | | 44.14 | 0.05 | 46.3 | 0.2 |
| NV6+MV5 | | | | | | | 47.36 | 0.04 | 49.84 | 0.02 |
| NV6+PV6 | | | | | | | 44.8 | 0.1 | 46.6 | 0.1 |
